# Conformational switching and flexibility in cobalamin-dependent methionine synthase studied by small-angle X-ray scattering and cryo-electron microscopy

**DOI:** 10.1101/2023.02.11.528079

**Authors:** Maxwell B. Watkins, Haoyue Wang, Audrey Burnim, Nozomi Ando

**Affiliations:** Department of Chemistry, Princeton University, Princeton, NJ, 08544, USA; Department of Chemistry and Chemical Biology, Cornell University, Ithaca, NY 14853, USA; Field of Biophysics, Cornell University, Ithaca, NY 14853, USA

## Abstract

Cobalamin-dependent methionine synthase (MetH) catalyzes the synthesis of methionine from homocysteine and 5-methyltetrahydrofolate (CH_3_-H_4_folate) using the unique chemistry of its cofactor. In doing so, MetH links the cycling of *S*-adenosylmethionine with the folate cycle in one-carbon metabolism. Extensive biochemical and structural studies on *Escherichia coli* MetH have shown that this flexible, multi-domain enzyme adopts two major conformations to prevent a futile cycle of methionine production and consumption. However, as MetH is highly dynamic as well as both a photosensitive and oxygen-sensitive metalloenzyme, it poses special challenges for structural studies, and existing structures have necessarily come from a “divide and conquer” approach. In this study, we investigate *E. coli* MetH and a thermophilic homolog from *Thermus filiformis* using small-angle X-ray scattering (SAXS), single-particle cryo-electron microscopy (cryo-EM), and extensive analysis of the AlphaFold2 database to present the first structural description of MetH in its entirety. Using SAXS, we describe a common resting-state conformation shared by both active and inactive oxidation states of MetH and the roles of CH_3_-H_4_folate and flavodoxin in initiating turnover and reactivation. By combining SAXS with a 3.6-Å cryo-EM structure of the *T. filiformis* MetH, we show that the resting-state conformation consists of a stable arrangement of the catalytic domains that is linked to a highly mobile reactivation domain. Finally, by combining AlphaFold2-guided sequence analysis and our experimental findings, we propose a general model for functional switching in MetH.

## Introduction

Modular, multi-domain enzymes catalyze remarkable chemical transformations by moving either their co-factor or substrate over large distances, from active site to active site. A classic example is the cobalamin-dependent methionine synthase (MetH) (Figure 1A), which is responsible for catalyzing methyl transfer from 5-methyltetrahydrofolate (CH_3_-H_4_folate) to L-homocysteine (Hcy) to produce H_4_folate and L-methionine (Figure 1B, black arrows). In humans, MetH activity is important for preventing elevated homocysteine levels, which are linked to increased risk of cardiovascular diseases and neural tube defects during embryonic development^1–6^. Additionally, MetH activity is essential for regenerating H_4_folate for one-carbon metabolism^1,7–11^, which supports a number of critical processes, such as nucleotide biosynthesis in dividing cells^11^.

**Figure 1:**
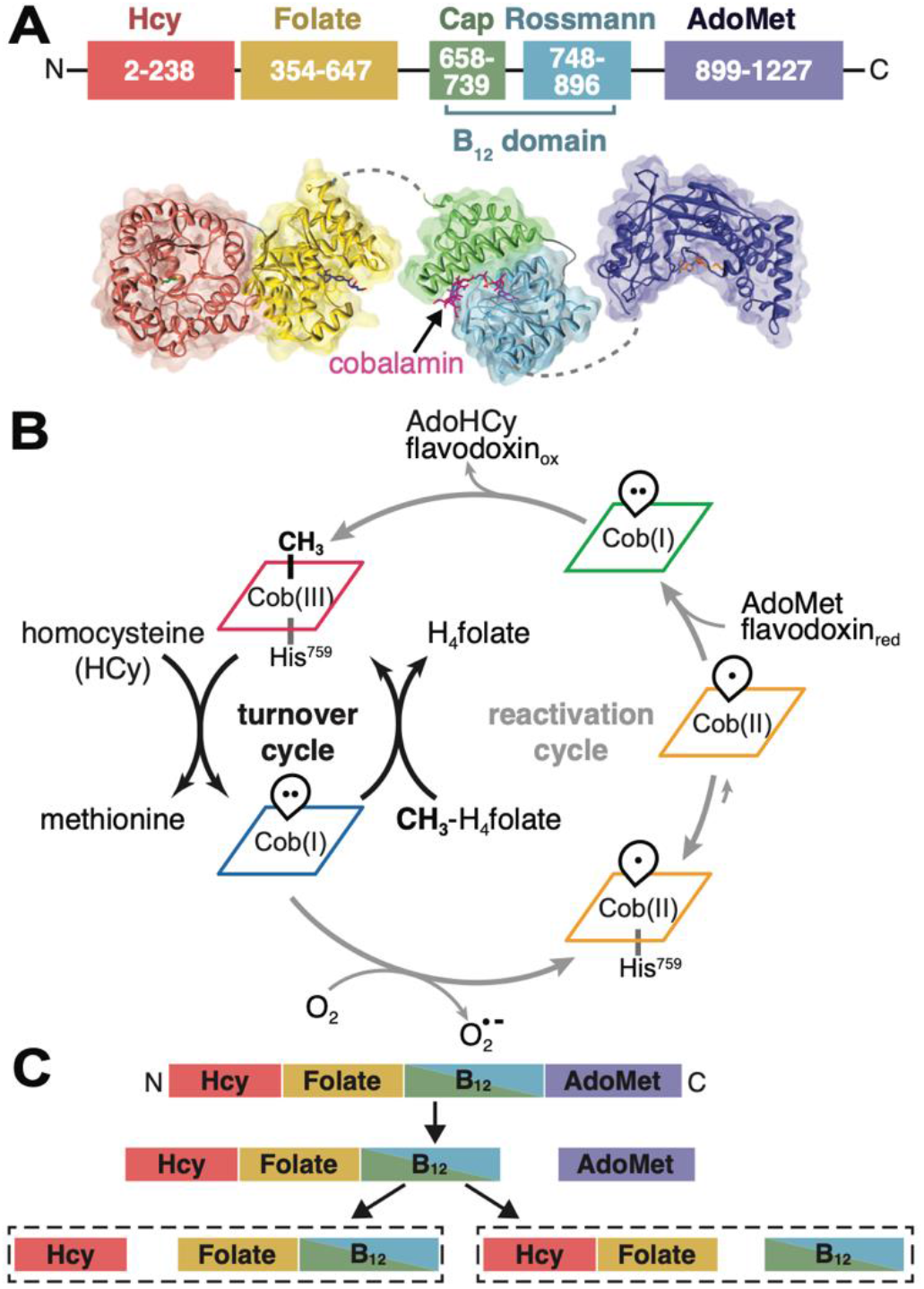
Overview of MetH. **(A)** MetH has a multi-domain architecture (shown in *E. coli* numbering), consisting of four domains, which bind homoycteine (Hcy), 5-methyltetrahydrofolate (CH_3_H_4_folate)^13^, cobalamin (B_12_)^17^ and *S*-adenosylmethionine (AdoMet)^19^. The B_12_ domain additionally consists of the cap and Rossmann subdomains. There is flexibility expected between all the domains except the two N-terminal domains. **(B)** During normal turnover (black arrows), CH_3_-Cob(III) enzyme will bind Hcy and CH_3_-H_4_folate sequentially, with the latter preferentially binding first^20^. Both reactions then proceed in the ternary complex, with the first reaction being methylation of Hcy by CH_3_-Cob(III) to yield methionine and a reduced Cob(I) intermediate. Methyl transfer from CH_3_-H_4_folate to the cofactor then yields H_4_folate and regenerates CH_3_-Cob(III). When the Cob(I) cofactor is occasionally oxidized to inactive Cob(II), MetH is able to reactivate via reductive methylation using AdoMet as a methyl donor and an electron from hydroquinone or semiquinone flavodoxin (grey arrows). Cob(II) is predominantly His-on at physiological pH but can interconvert with the His-off state, which is likely to be recognized by flavodoxin. The reactivation cycle proceeds through a distinct, 4-coordinate Cob(I), which is thought to be conformationally gated from normal turnover. **(C)** Limited trypsin proteolysis produces two different pathways. Cob(II) enzyme in the His-off state or in the presence of flavodoxin proceeds through the right pathway, while all other states proceed through the left^12^.

An extensive body of biochemical and structural work by Matthews, Ludwig, and co-workers has provided significant insight into MetH mechanism, especially for the *E. coli* enzyme^12–24^. The free form of MetH is thought to be in the His-on CH_3_-Cob(III) (methylcobalamin) oxidation state^20^ (Figure 1B, red), with a conserved histidine (His759 in *E. coli* numbering) acting as a lower ligand to the central cobalt of the cofactor. In this state, the enzyme is able to transfer the methyl group from its cofactor to Hcy, generating methionine and reducing the cofactor to the Cob(I) oxidation state (Figure 1B, blue), which is preferentially His-off. The CH_3_-Cob(III) form of the enzyme is then regenerated by methyl transfer from CH_3_-H_4_folate, producing H_4_folate. Under physiological conditions, MetH is inactivated by oxidation to a predominantly His-on Cob(II) state (Figure 1B, orange) approximately once every 2000 turnovers^12^. During the reactivation cycle (Figure 1B, grey arrows), the enzyme is able to re-activate through reductive methylation of the cofactor through electron transfer from reduced flavodoxin and methyl transfer from *S*-adenosylmethionine (AdoMet). Binding of flavodoxin favors the dissociation of the His759 ligand in the Cob(II) state (Figure 1B, orange/His-off) and the enzyme is able to reenter the turnover cycle via a distinct Cob(I) intermediate^15,21^ that is reactive with AdoMet but not with CH_3_-H_4_folate (Figure 1B, green).

Remarkably, the three reactions catalyzed by *E. coli* MetH imply significant conformational rearrangements of four distinct domains. From N-to C-terminus, these domains are known as the Hcy domain, folate domain, B_12_ domain, and AdoMet or reactivation domain^24^ (Figure 1A). Crystal structures of MetH fragments from various organisms have yielded important mechanistic insight^13,15–17,19,25^. The structure of the *E. coli* B_12_ domain captured two subdomains in a so-called “cap-on” conformation, with the cobalamin (Figure 1A, magenta) bound in the Rossmann-fold subdomain (Figure 1A, cyan) and the upper face covered by a four-helix bundle cap subdomain^17^ (Figure 1A, green). Structures of the two N-terminal domains from *Thermotoga maritima*^13,16^ MetH revealed a back-to-back double triose-phosphate isomerase (TIM)-barrel architecture (Figure 1A, salmon/yellow), which suggested that considerable domain motions in the rest of the enzyme are required to shuttle the cobalamin back and forth between the two active sites during turnover. A third active site was visualized in the C-shaped structure of the *E. coli* AdoMet domain (Figure 1A, navy). However, despite decades of extensive research, the full-length MetH remained elusive to structure determination because of its flexible “beads on a string” arrangement.

Seminal work by Jarrett, et al. have shown that there must be two major conformations of *E. coli* MetH that enable the enzyme to distinguish its two methyl donors, CH_3_-H_4_folate and AdoMet, and electron transfer partner, flavodoxin^12^. In this study, it was shown that tryptic proteolysis of *E. coli* MetH proceeds through two distinct pathways, depending on the cofactor oxidation state and the presence of flavodoxin. In both pathways, the 38-kDa AdoMet domain is the first to be cleaved (Figure 1C). Intriguingly, subsequent cleavage patterns for the two Cob(I) states, His-on Cob(II) state, and CH_3_-Cob(III) state were highly similar, with the Hcy domain separating from a fragment that contains both the folate and B_12_ domains (Figure 1C, left). In contrast, when Cob(II) enzyme was in the presence of flavodoxin or made to be constitutively His-off (via a H579G mutation), the B_12_ domain was removed from the two N-terminal domains (Figure 1C, right). Spectroscopic studies^26,27^ have shown that although free Cob(II) MetH is predominantly His-on at physiological pH, flavodoxin favors the formation of the His-off state and binds strongly to the H579G mutant. These observations indicated that that the second proteolytic pathway must represent a conformation of MetH is recognized and/or stabilized by flavodoxin and therefore important for reactivation of the cofactor. This so-called “reactivation conformation” was later established by crystal structures of the two C-terminal domains of *E. coli* MetH, in which a cap-off B_12_ domain is captured interacting with the active site of the AdoMet domain^14,15,18^. Because the two N-terminal domains are presumably uninvolved in this conformation, it would make sense that they are cleaved as a single unit when subjected to proteolysis (Figure 1C, right). Together, these findings implied that conformational switching in MetH is governed by the oxidation state of the cobalamin cofactor in some way. Since AdoMet is derived from methionine, such a conformationally gated mechanism would prevent a futile cycle of methionine production and consumption.

In this study, we used a combination of advanced small-angle X-ray scattering (SAXS), cryo-electron microscopy (cryo-EM), and extensive exploration of the AlphaFold2 database to report the first structural description of full-length MetH and the mechanism by which the enzyme switches functional modes. We show with SAXS that *E. coli* MetH shares a conformational state in both active and inactive oxidation states and that CH_3_-H_4_folate is uniquely able to drive a conformational change even when it is not a reactant. We further show with cryo-EM and SAXS that this shared conformational state is one in which a cap-on B_12_ domain is nested behind the two N-terminal domains in a “resting state” and the C-terminal AdoMet domain undergoes continuous motion. The cryo-EM model, derived from a thermostable MetH from *T. filiformis*, provides an explanation for why CH_3_-H_4_folate binding undocks the B_12_ domain, leading to a conformation predicted by SAXS and AlphaFold2, where a cap-off B_12_ domain interacts with the folate domain. In parallel, we show by SAXS that although the different oxidation states are conformationally indistinguishable, flavodoxin preferentially binds the Cob(II) state. With these results, we propose a model for MetH structural dynamics that is consistent with key biochemical findings. In this model, a stable resting-state conformation is shared between the turnover and reactivation cycles and the presence of CH_3_-H_4_folate triggers a conformational change that initiates turnover, while recognition by flavodoxin recruits Cob(II) MetH into the reactivation cycle.

## Results

### MetH oxidation states share a conformation in the absence of substrates

Based on prior studies, the full-length *E. coli* MetH was expected to be highly flexible. We therefore began by comprehensively characterizing this enzyme using SAXS. SAXS is unique in its ability to yield direct structural information on an entire solution conformational ensemble and, when applied carefully, can detect subtle conformational changes in flexible, multi-domain proteins^28,29^. To determine how the cobalamin oxidation state correlates with conformation, we compared three representative states of MetH: the CH_3_-Cob(III) and Cob(I) states from the turnover cycle and the inactive, His-on Cob(II) state from the reactivation cycle (Figure 1B). In wild-type MetH, the CH_3_-Cob(III) cofactor is known to be almost entirely His-on over a wide range of solution conditions, while the His-on/off equilibrium in the Cob(II) cofactor is dependent on pH, with the His-on state being predominant at physiological pH^26,30^.

We first performed size-exclusion chromatography-coupled SAXS (SEC-SAXS) to determine the number of distinguishable species in the absence of substrates. *E. coli* MetH is known to purify as a mixture of physiological and non-physiological cobalamin oxidation states, from which each physiologically relevant state shown in Figure 1B can be prepared enzymatically, chemically, or via photolysis^31^. SEC-SAXS of as-isolated MetH produces a single elution peak (Figure S1A). However, singular value decomposition (SVD) of the SEC-SAXS dataset reveals that the peak contains at least three components (Figure S1A), two of which are highly overlapped near the center of the peak (Figure S1B). In contrast, MetH prepared in a single oxidation state predominantly elutes as a single species (Figures S1C-D and S2A-D). Together, these results indicate that purifying the oxidation state of MetH also removes the heterogeneity in conformational state that is seen in the as-isolated enzyme.

Having established by SEC-SAXS that MetH prepared in an individual oxidation state cannot be further separated by chromatography (Figures S2C-D), we switched to batch-mode SAXS experiments, which yielded highly similar profiles (Figure S2E-F). Batch mode was ultimately necessary because certain substrates of MetH are not readily available in large enough quantities for use in the running buffer in SEC-SAXS experiments. Additionally, these experiments offer greater control over verifying the integrity of the sample oxidation state. To gain insight into structural differences between oxidation states, we examined the representative states of the full-length *E. coli* MetH in the absence of substrates in reducing buffer (50 mM HEPES pH 7.6, 150 mM NaCl, 2.5 mM DTT). UV-Vis absorption spectra were collected on each sample before and after X-ray exposure to monitor any changes in cobalamin oxidation state. In reducing buffer, His-on Cob(II) MetH is stable and amenable to standard experimental setups. The integrity of the 5-coordinate, His-on Cob(II) state was verified by the presence of a peak at ~477 nm^27,31^ both before (Figure 2A, orange) and after (Figure S3B, inset) X-ray exposure. CH_3_-Cob(III) MetH, on the other hand, is photosensitive^23^ and necessitated a fully darkened experimental hutch, with illumination sources limited to red light. The observation of a broad peak at ~528 nm^27,31^ both before (Figure 2A, red) and after (Figure S3C, inset) exposure was an indication that the 6-coordinate, His-on CH_3_-Cob(III) state did not undergo photolysis or photoreduction. Finally, a fully anoxic setup was required for the highly oxygen-sensitive Cob(I) MetH^20,31^, where the sample loading, pumps and waste lines were contained in an in-line anoxic chamber at the beamline (SI Methods). The oxidation state was verified by the characteristic, sharp peak at ~390 nm^27,31^ observed both before (Figure 2A, blue) and after (Figure S3A, inset) X-ray exposure.

**Figure 2:**
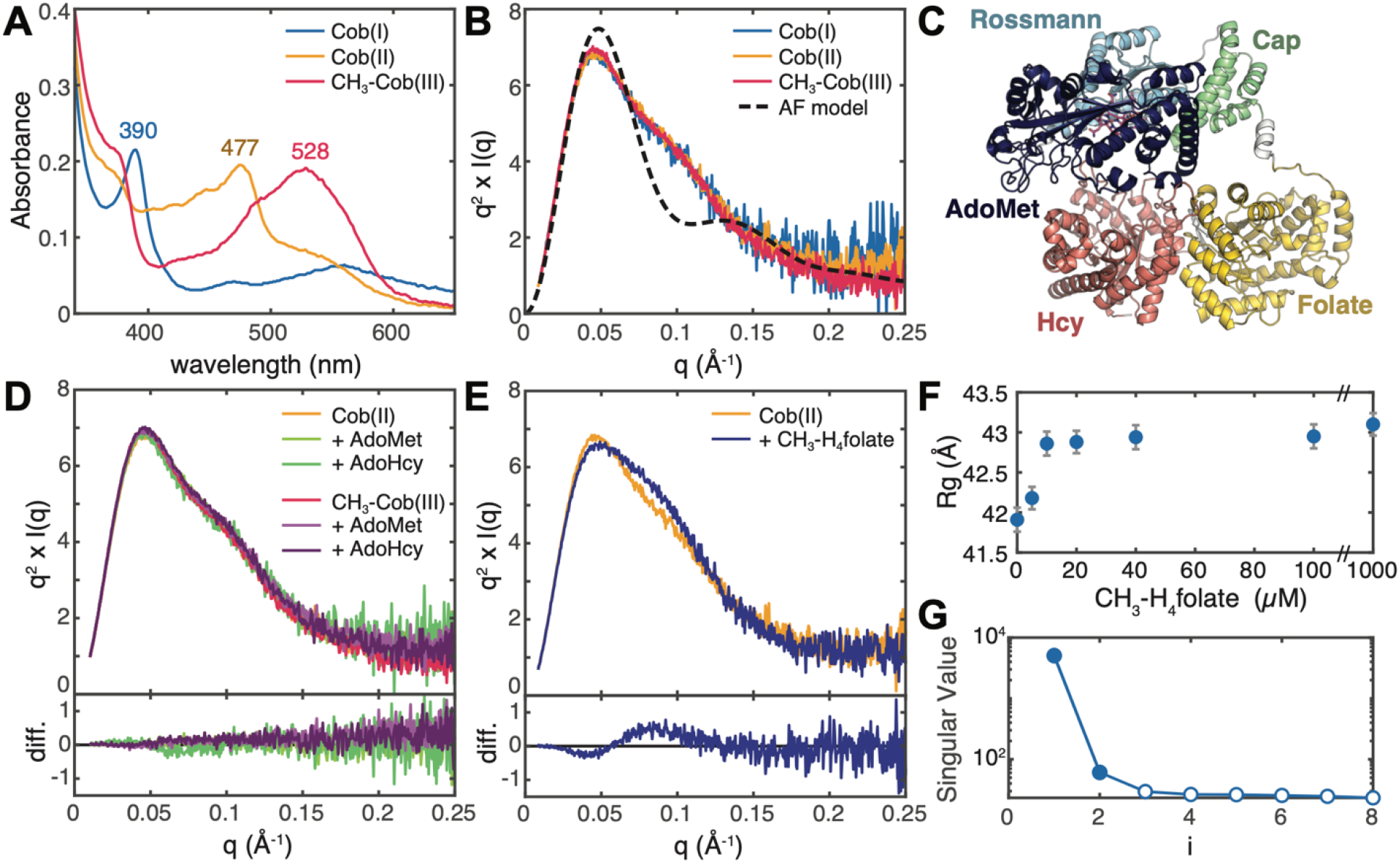
SAXS shows that there is a shared conformation of *E. coli* MetH and that conformational change is driven by CH_3_H_4_-folate. **(A)** UV-Vis absorption spectra of *E. coli* MetH in the Cob(I) and CH_3_-Cob(III) states from the turnover cycle (blue and red) and the inactive Cob(II) state (orange) are distinct. **(B)** SAXS profiles, shown in Kratky representation (*q*^2^ x *I* vs. *q*), of 20 μM *E. coli* MetH in the three oxidation states (same colors as in panel A) are indistinguishable. The theoretical scattering from the AlphaFold2 model in panel C (black dashed) displays a much sharper Kratky peak compared to the experimental curves, indicating that in solution MetH is much more extended and flexible. **(C)** The AlphaFold2 prediction for full-length *E. coli* MetH predicts the two C-terminal domains in the reactivation conformation. **(D)** Addition of reactivation-cycle substrates and products produces no discernable change to the MetH conformation. Top plot shows the Kratky curves, while the bottom shows difference curves. **(E)** Addition of 1 mM CH_3_H_4_-folate to Cob(II) MetH results in a widened Kratky peak, indicative of a conformational change to a more elongated structure. Top plot shows the Kratky curves, while the bottom shows difference curves. **(F)**The conformational change corresponds to a modest increase in the radius of gyration (*R_g_*) with increasing CH_3_*-*H_4_folate concentration. The error bars represent the standard deviation from Guinier analysis. **(G)**SVD of the titration series shown in panel F suggests a minimally two-state transition.

SAXS profiles were obtained on 20 μM *E. coli* MetH in the three oxidation states. To facilitate detection of any subtle differences in the enzyme domain arrangement, the scattering is shown in Figure 2B in Kratky representation: *Iq*^2^ vs *q*, where *I* is the scattering intensity, *q*=4π/*λ* sin*θ* is the momentum transfer variable, 2*θ* is the scattering angle, and *λ* is the X-ray wavelength. The Kratky plots for all three states are essentially indistinguishable (Figure 2B, solid), exhibiting a skewed peak with a maximum at *q*~0.045 Å^-1^ with a characteristic subtle shoulder at *q*~0.11 Å^-1^ that slowly decays at higher *q*. In comparison, the theoretical Kratky plot of an AlphaFold2^32^ model of the full-length *E. coli* MetH (Figure 2C) displays a much sharper peak at *q*~0.05 Å^-1^ (Figure 2B, dashed). The slow decay at high *q* seen in the experimental Kratky plots indicates that in solution, MetH samples conformations that are more extended and flexible than that captured by the AlphaFold2 model. Consistent with this, the experimental *R_g_* values obtained by Guinier analysis range between ~40-42 Å (Table S2), which is significantly larger than the theoretical *R_g_* value of 35.2 Å for the AlphaFold2 model. These larger *R_g_* values are not due to oligomerization. Based on Porod volumes (*V_P_*) calculated from the SAXS profiles, the molecular weights of the CH_3_-Cob(III), Cob(II), and Cob(I) states are estimated to be 138.4, 138.9, and 142.1 kDa, respectively, consistent with monomeric MetH (actual molecular weight of holoenzyme = 137.2 kDa). We note that Cob(II) and Cob(I) MetH have slightly higher apparent *R_g_* values and maximum dimensions (*D_max_*) than CH_3_-Cob(III) MetH (Table S2, Figure S3). These small differences are not particularly meaningful as the SAXS profiles overlay almost exactly and are instead attributed to small amounts of additional aggregate resulting from the chemical preparation of those states from the CH_3_-Cob(III) enzyme. Most importantly, these results suggest that MetH is flexible in solution and that in the absence of substrates, the different oxidation states are likely to share a conformation or set of conformations. This observation explains for the first time why these oxidation states share a proteolytic pathway (Figure 1C, left)^12^.

### Conformational changes are driven by CH_3_H_4_folate

It was previously shown by UV-Vis absorption spectroscopy that the interconversion of His-on and His-off states of the cobalamin cofactor can be modulated by substrates and products in a mutant of *E. coli* MetH where the His-on state is destabilized^33^. To test whether these substrates and products can shift the conformational distribution of wild-type *E. coli* MetH, we examined two representative states of the turnover and reactivation cycles, namely the His-on Cob(II) and CH_3_-Cob(III) states, in reducing buffer (50 mM HEPES pH 7.6, 150 mM NaCl, 2.5 mM DTT) using batch-mode SAXS. A maximum substrate concentration of 1 mM was chosen to match previously reported experiments and to saturate binding based on known dissociation constants^12,20,33^ (SI Methods).

We first examined the effects of the substrate and product pair, *S*-adenosyl-L-methionine (AdoMet) and *S*-adenosylhomocysteine (AdoHcy), from the reactivation cycle. Incubation of 20 μM CH_3_-Cob(III) or Cob(II) *E. coli* MetH with either 1 mM AdoMet or AdoHcy produced no appreciable change in the SAXS profiles relative to the substrate-free enzyme (Figures 2D, S4A-B). Likewise, we examined the substrate and product pair, homocysteine (Hcy) and methionine (Met), from the turnover cycle. Incubation of Cob(II) enzyme with either 1 mM Met or Hcy again led to identical profiles as the substrate-free enzyme (Figure S4C). The CH_3_-Cob(III) enzyme was tested only in the presence of 1 mM Met as Hcy would rapidly react with the enzyme, but again, no change in scattering was observed (Figure S4D).

Interestingly, when Cob(II) enzyme was incubated with CH_3_-H_4_folate (specifically, (6*S*)-5-CH_3_H_4_-PteGlu3), a clear change in scattering was observed (Figure 2E). Titration of 0-1 mM CH_3_-H_4_folate into 20 μM Cob(II) MetH led to the appearance of a shoulder on the Kratky peak at *q* ~ 0.09 Å^-1^ (Figures 2E, S5A), indicative of a conformational rearrangement that is more elongated than the substrate-free enzyme. Consistent with this interpretation, there is a shift in the pair distance distribution, *P*(*r*), towards a shape that is more consistent with an elongated species (Figure S5B). This change was accompanied by a modest increase in the Guinier *R_g_*, from 41.9 ± 0.2 Å in the absence of substrates to 43.1 ± 0.1 Å after incubation with 1 mM CH_3_-H_4_folate (Figures 2F, S5D, Table S3). SVD of the titration series suggested a minimally two-state transition (Figure 2G). Addition of CH_3_-H_4_folate to CH_3_-Cob(III) MetH resulted in an identical change (Figure S5C), although the enzyme was unavoidably reduced by the X-ray beam to Cob(II) enzyme during the experiment (Figure S5C, inset), and thus the resulting scattering likely represents that of a mixture of oxidation states, if not of pure Cob(II) enzyme. Nonetheless, these results are significant for several reasons. Of the many conditions we tested, CH_3_-H_4_folate was the only ligand that induced a detectable conformational change, despite the fact that it is not a substrate for either the Cob(II) or CH_3_-Cob(III) enzymes. This suggests that CH_3_-H_4_folate is generally responsible for driving conformational change in MetH. Such an interpretation is in line with previous steady-state kinetics results that show that the primary turnover reaction involves an ordered sequential mechanism, which is driven by CH_3_-H_4_folate preferentially binding first in the ternary complex^20^. Furthermore, we found that CH_3_-Cob(III) MetH reduces to Cob(II) in the X-ray beam when CH_3_-H_4_folate is bound but not in the absence of substrates or in the presence of AdoMet, AdoHcy, or Met. This result is consistent with previous studies that observed enhanced photolysis rates of the CH_3_-Cob(III) cofactor upon addition of CH_3_-H_4_folate^23^ and suggests that the conformational change induced by this substrate involves the uncapping of the B_12_ cap domain, which normally acts to sequester the methyl radical produced by the homolysis of the methylcobalamin Co-C bond. This, in turn, indicates that there exists a resting state of MetH, in which the B_12_ domain is capped.

### Flavodoxin binding is specific to the cobalamin oxidation state

Our SAXS data thus far suggest that both turnover and reactivation states of MetH share a so-called “capon” resting-state conformation that undergoes a similar response to CH_3_-H_4_folate. To gain insight into what initiates the reactivation cycle, we examined complex formation between MetH and flavodoxin using SAXS. Although flavodoxin binding to the Cob(II) enzyme is relatively weak at near physiological pH (*K_D_* ~ 46.5 μM at pH 7.0), extrapolation of pH-dependent data^26^ suggests that complex formation should be readily discernable by SAXS around pH 6 at MetH concentrations consistent with our other experiments.

Exchanging CH_3_-Cob(III) and Cob(II) MetH into reducing buffer at pH 6 (44 mM Na/K phosphate pH 6.0, 1 mM DTT with the ionic strength adjusted to 0.15) leads to SAXS profiles that are highly similar to those obtained at pH 7.6 (Figure S6A-B). The subtle pH-dependent change in Cob(II) scattering at *q*~0.045 Å^-1^ (Figure S6B, arrow) may reflect a slight increase in His-off population (which has an absorbance peak at 465 nm)^26^ at pH 6, as indicated by the slight shift in the absorbance peak from 477 to 472 nm (Figure S6C). Despite minor differences in the scattering of free CH_3_-Cob(III) and Cob(II) MetH, markedly different behavior is observed when the enzyme is mixed with flavodoxin (Figure 3A,D). Addition of 0-60 μM *E. coli* flavodoxin to 20 μM Cob(II) MetH in reducing buffer leads to an increase in the apparent Guinier *R_g_* value until approximately the equimolar point, followed by a decrease in *R_g_* (Figure 3B, Table S4). The initial increase in *R_g_* is consistent with complex formation, while the subsequent reduction in apparent *R_g_* can be explained by excess amounts of the titrated protein. In agreement with this interpretation, SVD of the titration series (Figure 3C) indicates a minimally three-component mixture, likely corresponding to free MetH, MetH:flavodoxin complex, and free flavodoxin. By contrast, when flavodoxin is titrated into 20 μM CH_3_-Cob(III) MetH, the observed behavior is a steady decrease in the Guinier *R_g_* value (Figure 3E, Table S5). SVD of the titration series suggests a minimally two-component system (Figure 3F), which when taken with the steady decrease in *R_g_*, indicates a lack of complex formation.

**Figure 3:**
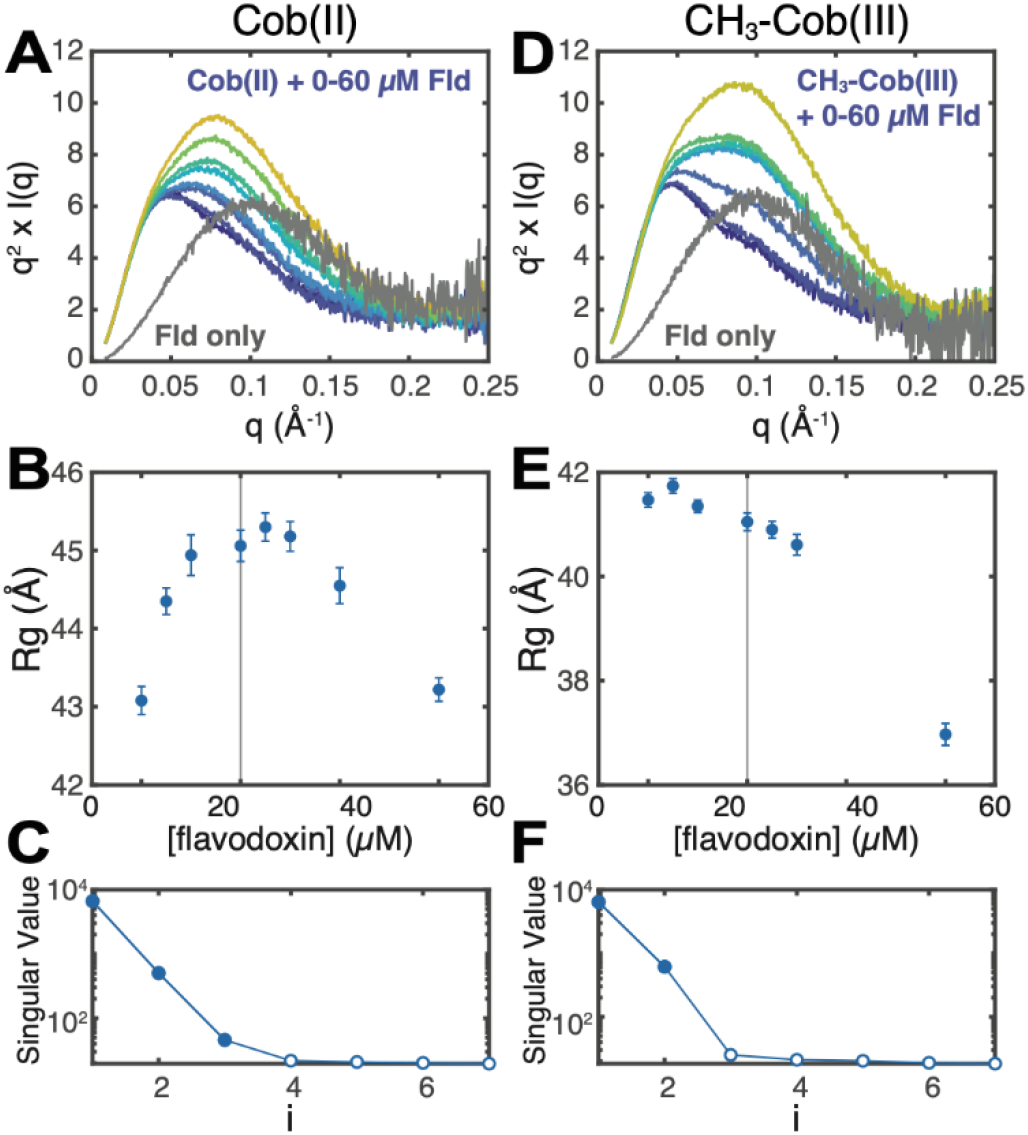
SAXS shows that flavodoxin discriminates between *E. coli* MetH oxidation states. **(A-C)** Titration of 0-60 μM *E. coli* flavodoxin (Fld) into 20 μM Cob(II) *E. coli* MetH at pH 6.0 produces behavior consistent with complex formation and saturation. **(A)** Kratky curves for the titration series are shown in color (navy, 0 μM Fld, to yellow, 60 μM Fld), while that of pure Fld is shown in grey. **(B)** The apparent Guinier *R_g_* of the mixture increases to near the equimolar point (vertical line), indicative of complex formation, followed by a decrease due to excess Fld (the *R_g_* of free Fld is ~18 Å). **(C)** SVD of the titration series shown in panel Fld suggests a minimally three-component system, consistent with the two proteins as well as a complex of the two. **(D-F)** By contrast, titration of 0-60 μM Fld into 20 μM CH_3_-Cob(III) MetH indicates a lack of complex formation. **(D)** Kratky curves for the titration series are shown in color (navy to yellow), while that of pure Fld is shown in grey. **(E)** The apparent Guinier *R_g_* of the mixture decreases with increasing Fld, indicating that the two proteins do not interact. **(F)** SVD of the titration series shown in panel D suggest a minimally two-component system, consistent a mixture of two non-interacting proteins.

These SAXS results clearly demonstrate that flavodoxin binds specifically to Cob(II) MetH but not CH_3_-Cob(III) MetH. In our experiment, both enzyme states were predominantly His-on before addition of flavodoxin. It is known, however, from spectroscopy and crystallography that binding of flavodoxin to Cob(II) MetH promotes the lower His759 ligand of the cofactor to dissociate^12,26^, and that in the reactivation conformation, His759 instead interacts with Glu1069 and Asp1093 in the AdoMet domain, stabilizing the interaction of the two C-terminal domains^18^. Combined with these previous observations, our results indicate that flavodoxin recognizes the oxidation state of MetH in which His759 readily dissociates from the cobalamin cofactor, which is the case for Cob(II) but not CH_3_-Cob(III). This in turn suggests that although the turnover and reactivation cycles share a resting-state conformation, the Cob(II) enzyme can enter the reactivation cycle through recognition by flavodoxin, which is always present *in vivo*.

### Cryo-EM structure of methionine synthase represents a resting state

To gain higher resolution insight, cryo-EM experiments with His-on Cob(II) and CH_3_-Cob(III) *E. coli* MetH were performed extensively both in the absence and presence of CH_3_-H_4_folate. However, these experiments were continually hampered by the destabilization environment of the air-water interface^34^, likely due to the well-known instability of the B_12_ domain in the *E. coli* enzyme^31^. To overcome these issues, we calculated a sequence similarity network (SSN) of 8,047 MetH sequences and identified several homologous sequences from thermophiles both within the same SSN cluster as the *E. coli* enzyme and elsewhere (Figure S7). Of the three sequences that we tested, we found that *Thermus filiformis* MetH (33% sequence identity, 53% sequence similarity to *E. coli*) was stable enough that it could be purified in the apo form and later be reconstituted with the cobalamin cofactor (SI Methods), suggestive of a more stable B_12_ domain fold than the *E. coli* enzyme. SAXS profiles of 20 μM Cob(II) *T. filiformis* MetH in the absence and presence of 1 mM CH_3_-H_4_folate are superimposable with the *E. coli* counterparts (Figure S8), indicating that the two enzymes behave similarly. Single-particle cryo-EM was then performed (Figure S9-10, Tables S7-8). To address severe orientation bias seen in screening datasets and to stabilize very thin ice at the hole centers, *T. filiformis* MetH was vitrified concurrently with equimolar horse spleen apoferritin, which ameliorated both issues. As apoferritin is comparable in size to MetH (as suggested by SAXS *D_max_*) and is known to rapidly denature at the air-water interface^35^, we attribute this improvement to apoferritin creating sacrificial denatured layers at the interfaces and potentially also acting as a physical support. We note however that addition of apoferritin did not help with cryo-EM attempts of *E. coli* MetH, and thus, the intrinsic stability of the *T. filiformis* enzyme appears to be the most important factor.

With this strategy, we ultimately obtained an accurate view of the intact, full-length *T. filiformis* MetH in the substrate-free Cob(II) state. 2D classes revealed clear secondary structure elements for the three N-terminal domains and blurry contrast for the C-terminal AdoMet domain (Figure 4A). These data gave rise to a 3.6-Å resolution consensus map of the three N-terminal domains (Figure 4B-C), while additional analyses (discussed in detail in the next section) demonstrated that the C-terminal AdoMet domain is highly dynamic in this state (Figure 5). The consensus map reconstruction represents the first structure of more than two domains of MetH in any state—a description that has remained elusive despite almost three decades of structural studies. Although the map exhibits some resolution anisotropy (Figure S10D), the overall resolution and detail of the map allowed for unambiguous placement of the individual protein domains as well as the cobalamin cofactor and the catalytic zinc ion in the Hcy-binding domain (Table S8). Overall, the sub-conformation of the two N-terminal domains is similar to previous fragment structures of *Thermatoga marihma*^13,16^ and human MetH (unpublished, PDB 4CCZ) (Figure S11A, bottom), where the N-terminal domains appear to adopt a rigid platform with the two active sites pointing away from each other. Additionally, the B_12_ domain adopts a highly similar overall conformation to the published fragment structure of the *E. coli* B_12_ domain (PDB 1BMT^17^, Figure S11A, top). In this conformation, the four-helix bundle cap subdomain sits atop the cobalamin cofactor, likely serving to protect it from solvent and against unwanted reactivity. Moreover, our cryo-EM map supports a conformation in which His759 is stabilized by Asp757 to axially coordinate to the cobalt of the cobalamin cofactor, much like that observed in the previous fragment structure^17^ (Figure S12). It is notable that a Cob(II) MetH gave an identical B_12_ domain conformation to the crystal structure of the *E. coli* fragment^17^, which contained the CH_3_-Cob(III) form of the cofactor. This observation is consistent with our SAXS studies which suggest that the different oxidation states of MetH share a conformational state.

**Figure 4:**
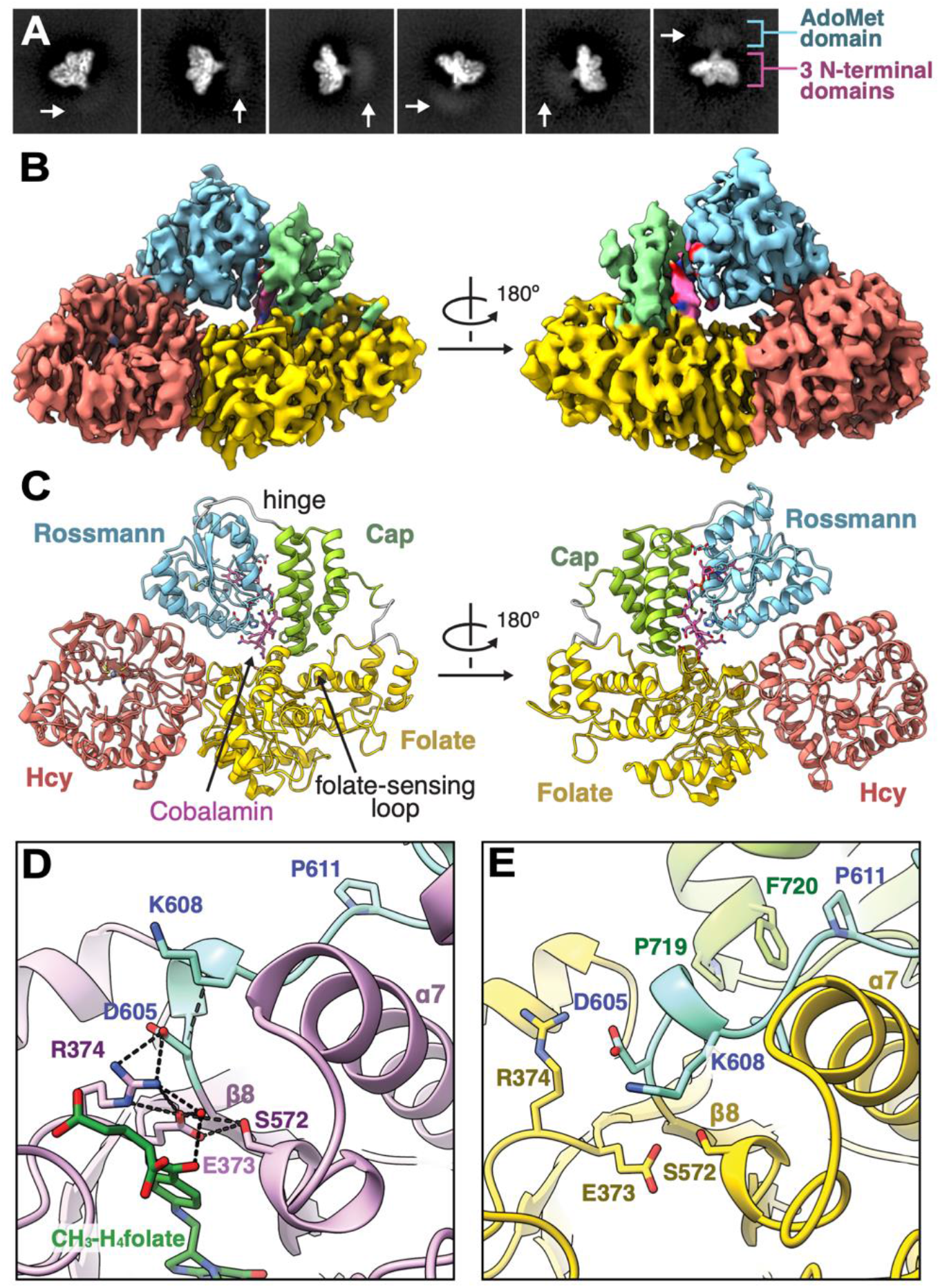
Cryo-EM of *T. Filiformis* MetH reveals a resting-state conformation poised for folate sensing. **(A)** 2D classes of Cob(II) *T. filiformis* MetH depicts the full-length enzyme. **(B)** The 3.6-Å consensus reconstruction contains the three N-terminal domains: the Hcy, folate, and B_12_ domains. The map is colored by domain assignment and shown at a threshold level of 0.3. **(C)** Residues 23-877 along with the cobalamin and zinc cofactors were built into the consensus map. The model suggests this conformation represents a resting state of the enzyme, where the B_12_ domain is in the cap-on conformation and nested between the Hcy and folate domains. **(D)** In a fragment structure of the folate domain from *T. thermophilus* MetH (PDB 5VOO)^25^, residues on the loop (cyan) that extends past the C-terminus of the folate-domain β8-strand (specifically, D605 and K608, shown in *T. filiformis* numbering for clarity) are oriented to help stabilize binding of CH_3_-H_4_folate. **(E)** In our cryo-EM structure, this folate-sensing loop (colored in cyan for contrast) is pushed inwards towards the folate-binding site by interactions with the cap subdomain (green). The competition between the cap subdomain and CH_3_-H_4_folate for the folate-sensing loop provides a structural basis for folate-driven disruption of the resting-state conformation and resulting conformational changes. Both hydrophobic and polar interactions are involved in stabilizing this closed conformation of the folate-binding loop (Figures S13-14).

Interestingly, the state captured by cryo-EM appears to represent a stable resting state of MetH. We observe the B_12_ domain tucked at the interface of the Hcy and folate domains (Figure 4C), such that the cofactor is sequestered from the active sites of both N-terminal domains. The cap subdomain of B_12_ domain interacts the folate domain through a combination of hydrophobic and polar contacts (Figure S13A-B, S14A), while the Rossmann subdomain binds the Hcy domain primarily through polar interactions (Figure S13C-D). The positioning of the two subdomains places N24 of the cobalamin cofactor within hydrogen-bonding distance of Thr366 on the folate domain. The interaction between the cap subdomain and folate domain is of particular interest because it involves a structural element that is not typically seen in a canonical β_8_α_8_ TIM barrel. In a canonical TIM barrel, the β8-strand connects via a loop to the α8-helix, which completes the barrel structure (Figure S15A). However, the MetH folate domain has been shown to adopt a β_8_α_7_ barrel topology, in which the loop from the β8-strand extends past the barrel wall and is terminated by a set of α8’-helices outside of the barrel (Figure S15B). Comparison of our cryo-EM structure with fragment structures of the folate domain^25^ reveals that this loop is in fact part of the active site (Figure 4D, blue loop) and that CH_3_-H_4_folate and the B_12_ cap subdomain must compete for interactions with this loop (Figure 4D-E, blue loop). Thus, the loop acts to sense the presence of CH_3_-H_4_folate, and binding of this substrate necessarily causes the B_12_ cap subdomain to undock from its resting-state position. In our structure, we observe hydrophobic interactions, including a favorable aromatic-proline stacking interaction between Phe720 on the cap domain and Pro611 in the folate-sensing loop, as well as polar interactions between Gln717 at the tip of the cap domain and Asp592 in the α7-helix of the folate domain (Figure S14A). In *E. coli* MetH, the positions of the Phe and Gln residues are flipped. Interestingly, analysis of the 4-domain MetH sequences in our SSN shows that majority can be classified into either *T. filiformis*-like or *E. coli*-like Q/F positioning in the cap domain. In *T. filiformis*-like sequences, Pro611 is highly conserved, while the residue at 592 is almost always one that can form a polar interaction with Gln (Figure S14B). Further sequence analysis and examination of other AlphaFold2 models in the resting-state conformation show that in *E. coli*-like sequences, Gln is positioned to form polar interactions with backbone of the folate-sensing loop, while the α7-helix of the folate domain contains highly conserved hydrophobic residues that can serve to conserve the overall interactions seen in the *T. filiformis* structure (Figure S14C-D). The conservation of overall interactions would explain why *T. filiformis* and *E. coli* MetH behave similarly in the absence and presence of CH_3_-H_4_folate. Together these interactions serve to “lock” the B_12_ domain in the cap-on conformation and sequester the cofactor from any reactivity.

The discovery of the resting-state conformation was unexpected, as interactions between the B_12_ domain and other MetH domains had not been predicted except during turnover or reactivation, when the cofactor must interact with the substrate-binding sites to facilitate methyl transfers. However, this resting-state conformation explains for the first time why limited proteolysis of MetH in any oxidation state first leads to a 98 kDa fragment containing the first three N-terminal domains^12^. Moreover, the interactions we observed between the cap subdomain and the folate domain explain why they remain associated in one of the proteolysis pathways^12^ (Figure 1C, left). Finally, our cryo-EM model explains how binding of CH_3_-H_4_folate can drive a conformational change from the resting-state conformation.

### Combining cryo-EM and SAXS yields a dynamic view of the resting state

Although the consensus map of the resting state lacks the AdoMet domain, blurry contrast is visible past the C-terminus of the B_12_ domain in the cryo-EM 2D classes (Figure 5A) that is suggestive of this domain undergoing significant motions. Consistent with this interpretation, swinging motion of the linker region is observed by 3D variability analysis in CryoSPARC^36^ (Movie S1). However, because the AdoMet domain is very small (~38 kDa) and mobile, it cannot be reconstructed by conventional methods.

**Figure 5:**
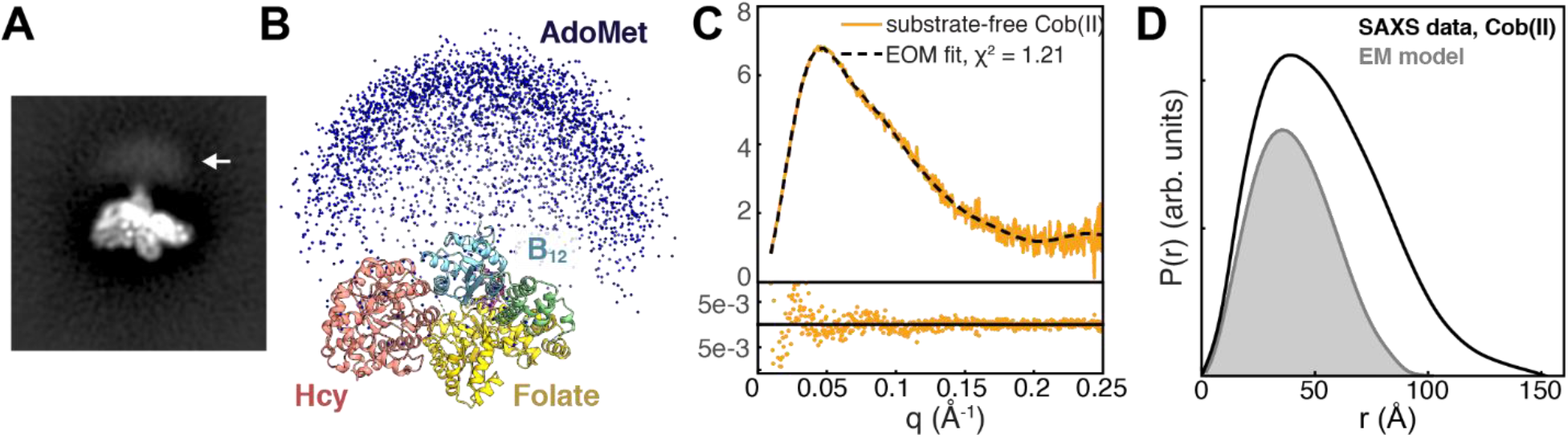
Combining cryo-EM with ensemble modeling of SAXS data shows that the AdoMet domain is highly flexible in the resting state of full-length MetH. **(A)** Blurry density is visible past the C-terminus of the B_12_-binding subdomain in reference-free 2D classes of *T. filiformis* MetH shown in Figure 4A (arrow). **(B)**A large ensemble of *E. coli* MetH structures was generated in EOM^37,38^ by treating the three N-terminal domains in the conformation observed by cryo-EM and the C-terminal AdoMet domain as two independent, flexibly-linked rigid bodies. Plotting the position of the AdoMet domain in each of the 10,000 models (blue spheres) produces a qualitatively similar distribution to the blurry density seen in the 2D classes in panel A. **(C)**The substrate-free Cob(II) *E. coli* MetH SAXS data from Figure 2B (orange) can be described very well by a minimal ensemble selected by EOM^37,38^ (dashed). **(D)**Comparison of the pair distance distribution, *P*(*r*), calculated from the SAXS data shown in panel C (black) to the theoretical *P(r*) generated from an *E. coli* model of the three N-terminal domains derived from the cryo-EM structure (grey) indicates that many of the large interparticle distances are not fully described by the EM model alone, indicating that the AdoMet domain samples many extended conformations. The *P*(*r*) functions were normalized by dividing by *I*(0)/MW^2^, where *I(0*) is the forward scattering intensity and MW is the molecular weight.

To probe the flexibility of the AdoMet domain, we combined the cryo-EM consensus model with SAXS data using a coarse-grained ensemble modeling approach known as ensemble optimization method (EOM)^37,38^. As much more is known biochemically about *E. coli* MetH than *T. filiformis* MetH, we chose to fit our models to the *E. coli* SAXS datasets. A resting-state model for *E. coli* MetH was built by aligning AlphaFold2 models for the three N-terminal domains to the cryo-EM consensus model of *T. filiformis* MetH. This model (consisting of residues 3-896) and a crystal structure of the *E. coli* AdoMet domain (residues 901-1227, from PDB 1MSK^19^) were treated as two independent rigid bodies connected by a 4-residue flexible linker. A coarse-grained ensemble of 10,000 structures sampling a wide range of conformational space was generated using the program RANCH^37,38^ (Figure 5B). A subset (i.e., a minimal ensemble) that provides the best fit to our experimental SAXS data was determined using a genetic algorithm implemented in the program GAJOE^37,38^. With this approach, we find that a minimal ensemble derived from the cryo-EM resting-state conformation and a flexibly linked AdoMet domain is able to describe our substrate-free Cob(II) SAXS data very well (Figure 5C).

The qualitative similarity of the 10,000 AdoMet domain positions in the starting EOM pool (Figure 5B) and the blurry contrast in the 2D classes (Figure 5A) suggests that the ensemble captured by cryo-EM samples an extremely wide range of conformational space. In EOM, the flexibility of the selected ensemble is assessed by *R_flex_*, which is 100% for a fully flexible system and 0% for a fully rigid system. By convention, the flexibility of the starting (i.e., random) ensemble is given in parentheses. For the EOM fit to the substrate-free Cob(II) SAXS data, *R_flex_* (random) is 84.80% (~ 89.73%), indicating that the selected ensemble represents a highly flexible system, having almost the same degree of the flexibility seen in the original structural pool. Additionally, comparing the experimental real-space SAXS distance distribution function, *P(r*), to one generated from the EM-based model (*E. coli* sequence) of the three N-terminal domains (Figure 5D) shows that a large fraction of the interparticle distances is not well described without the AdoMet domain. Taken together, these results are consistent with a resting-state model for the full-length MetH, in which the linker to the AdoMet domain is highly flexible, as suggested by CryoSPARC 3D variability analysis^36^ (Movie S1). That this domain could act almost as an independent protein is not surprising. In fact, in some species of bacteria including *T. maritima*, the AdoMet domain is a separate polypeptide chain entirely^39^, suggesting a minimal penalty for largely independent functions of the catalytic domains and the reactivation domain.

### A model for conformational change in MetH

As structure prediction continues to improve with the advent of AlphaFold2^32^ and RoseTTAFold^40^, our avenues for structural interpretation and validation continue to expand. In the case of MetH, the vast majority of public AlphaFold2 database models (2992 out of 4915 4-domain sequences) represent the reactivation conformation (Figures 2C, 6D), in which the B_12_ domain is uncapped and interacting closely with the AdoMet domain. This is not unexpected, as all available crystal structures of the two C-terminal domains of the enzyme adopt this conformation^14,15,18^, and machine learning enforces agreement with existing structures in the training dataset. Interestingly, however, the most recent public rank-0 model of the *T. filiformis* enzyme is strikingly similar to our observed cryo-EM resting-state model (Cα RMSD=2.5 Å for the three N-terminal domains, Figures 6A, S11B), although older versions of the database provided models in the reactivation conformation. In fact, running AlphaFold2 locally on the *T. filiformis* sequence without the AdoMet domain results in all 5 ranked models adopting a domain arrangement similar to the experimentally observed resting state. Moreover, of the 4915 MetH sequences in the public AlphaFold2 database with AdoMet domains, 212 are predicted to be in a resting-state conformation with the Rossmann domain within 2.5 Å Cα RMSD of the cap-on fragment structure (1BMT)^17^ (Figure S16A). This is an astonishing result, especially given that the prediction lacks the cobalamin cofactor.

**Figure 6:**
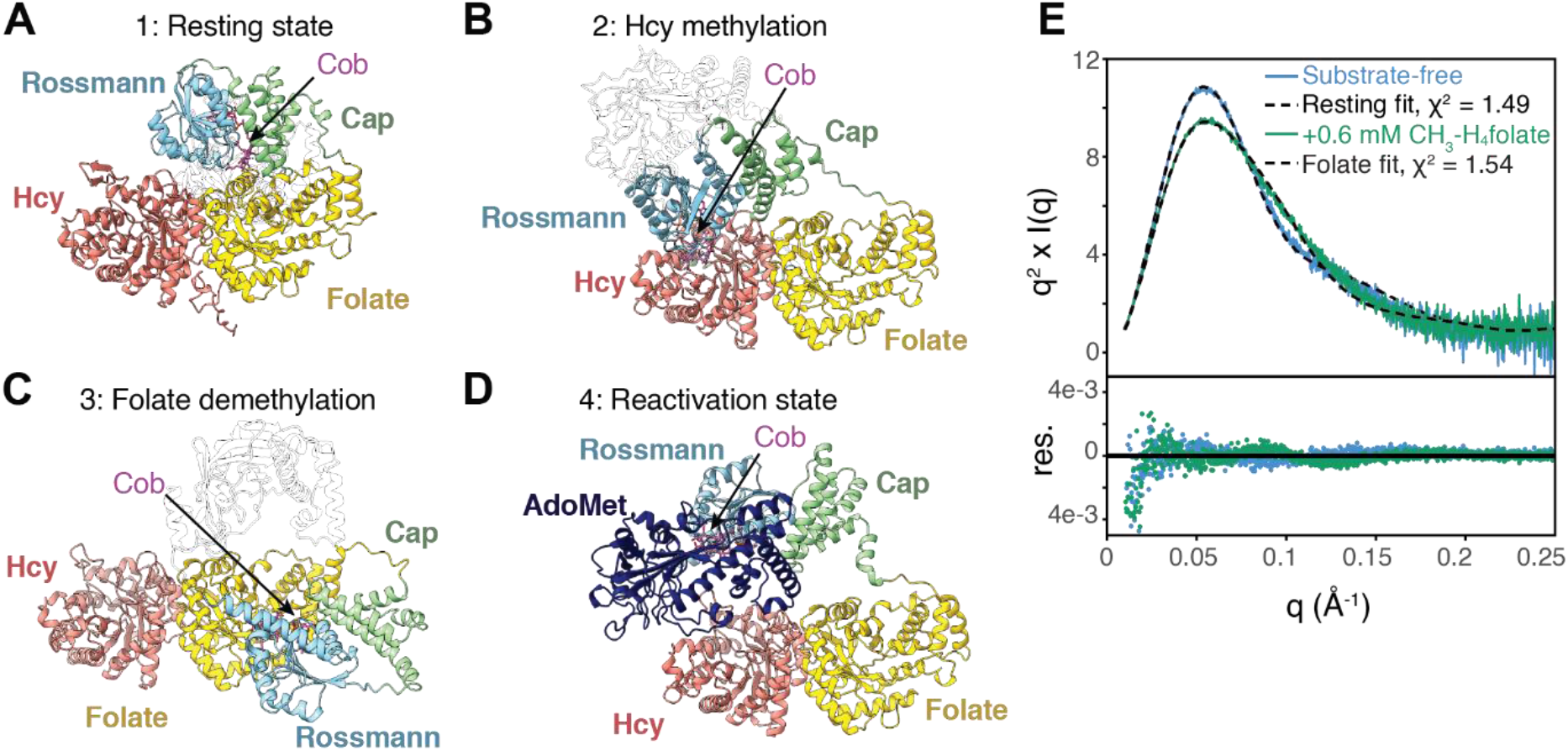
Structure prediction and removal of the AdoMet domain disambiguates the conformational change caused by CH_3_H_4_folate. **(A)** The most recent AlphaFold2 model for *T. filiformis* MetH predicts a conformation highly similar to the resting state observed in cryo-EM. **(B)** The AlphaFold2 model for *Mus musculus* MetH predicts a conformation similar to what would be expected to be competent for homocysteine methylation. **(C)** The AlphaFold2 model for a *Sphaerochaeta sp*. (Uniprot A0A521IQ66) predicts a conformation similar to what would be expected to be competent for folate demethylation. **(D)** The AlphaFold2 model for *E. coli* MetH predicts the C-terminal half of the enzyme in the reactivation conformation. This conformation is the most common conformation predicted among AlphaFold2 models, likely due to the prevalence of reactivation-state structures in the PDB. Models from across multiple species suggest that the N- and C-terminal halves do not interact. **(E)** A fragment containing only the three N-terminal domains of Cob(II) *E. coli* MetH was produced by tryptic proteolysis. The Kratky plot for this fragment (at 24 μM) displays a sharp peak (blue), indicating that removal of the AdoMet domain leads to a compact structure. Incubating the fragment with 0.6 mM CH_3_*-*H_4_folate leads to a change in scattering (green), indicating that folate-driven conformational change occurs within the three N-terminal domains. The model based on the cryo-EM resting state describes the substrate-free data well (blue/dashed), while a model based on that shown in panel C describes the folate-incubated Cob(II) data well (green/dashed). For panels A-C, the AdoMet domain has been rendered transparent for clarity, and for all models, the cobalamin ligand (magenta) has been docked into the structure by aligning the 1BMT^17^ structure to the cobalamin-binding loop of each model.

Based on the success of AlphaFold2 to predict a conformation that was observed experimentally for the first time in this study, we analyzed all 4915 publicly available models (SI Methods). Notably, a number of models appear to adopt a conformation similar to what would be expected during homocysteine methylation, i.e. with the binding face of the Rossmann subdomain situated above the barrel opening in the Hcy domain. These include models of the 3-domain *T. maritima* and 4-domain *Mus musculus* enzymes (Figure 6B). In the case of the *M. musculus* MetH model, cobalamin can be docked in without introducing serious steric clashes, placing the cofactor in a position that would put the cobalt in reasonable proximity to the homocysteine thiol. Of the 4915 publicly available AlphaFold2 models, 187 adopt a conformation in which the three N-terminal domains are within Cα RMSD of 2.5 Å of the *M. musculus* model (Figure S16B). Additionally, a handful of publicly available predicted models adopt a conformation similar to what would be expected during folate demethylation, i.e. with the binding face of the Rossmann subdomain situated above the barrel opening in the folate domain. These include the 4-domain MetH from a *Sphaerochaeta sp*. (Uniprot A0A521IQ66, Figure 6C) and 5 other models that are within 2.5 Å Cα RMSD (Figure S16C). The 1512 remaining models largely predict conformations in which the B_12_ domain is somewhere along a trajectory between the resting-state and the homocysteine-methylation positions (Figure S16A-B), possibly suggesting a path between these two states. Interestingly, these intermediate models do not align well with the cap-on B_12_ domain structure^17^ and therefore represent partially or fully uncapped conformations. In all predicted models not in the reactivation conformation, the AdoMet domain is randomly placed, consistent with the notion that it is flexibly linked.

To develop a model for conformational change in MetH without the ambiguity created by the flexible AdoMet domain, we performed limited proteolysis of the full-length Cob(II) *E. coli* MetH following established protocols^12^. A 98-kDa fragment containing only the three N-terminal domains of *E. coli* MetH (residues 2-896) was prepared to high purity and verified to be in the His-on Cob(II) state (Figure S17). SAXS was first performed on this fragment at 23 μM concentration in the absence of substrates. Remarkably, the Kratky plot displays a sharp peak at *q* ~ 0.055 Å^-1^ that decays quickly at high *q* (Figure 6E and S18, blue), indicating that the flexibility seen in the full-length enzyme is lost and that the three N-terminal domains must adopt a compact conformation in solution. In fact, the resting-state model of the three N-terminal domains from cryo-EM provides an excellent fit to the substrate-free SAXS data (Figure 6E, blue/dashed). Incubation of the 98 kDa fragment at 23 μM with 0.6 mM CH_3_-H_4_folate leads to a Kratky plot with a slightly broader peak (Figures 6E and S18, green), clearly indicating a substrate-induced conformational transition within the three N-terminal domains of MetH, from the resting state to one that is more extended in shape. Unexpectedly, we find that the elusive folate-demethylation conformation predicted by AlphaFold2 provides a good fit to the data (Figure 6E, green/dashed). We note that the AlphaFold2-predicted homocysteine-methylation conformation provides an equally good fit to the substrate-free data as the resting state (Figure S19). This is not too surprising as these conformations are highly similar at low resolution. However, based on our X-ray experiments of the full-length enzyme, which showed that CH_3_-Cob(III) is stable to photoreduction in the absence of substrates, we can gain confidence that the B_12_ domain is primarily capon in the free enzyme, which is the case for the resting-state conformation but not the homocysteine-methylation conformation.

## Discussion

Seminal work by Jarrett, et al. predicted the existence of two major MetH conformations that would enable the enzyme to distinguish its two methyl donors, CH_3_-H_4_folate and AdoMet^12^. Crystal structures of C-terminal fragments of *E. coli* MetH^14,15,18^ have shown that one of these major conformations must be that of the reactivation state, in which a cap-off B_12_ domain interacts with the AdoMet domain. In our study, we reported the existence of a resting-state conformation, in which the three N-terminal domains adopt a compact configuration that secures the B_12_ domain in a cap-on state, while the C-terminal AdoMet domain is highly mobile. Because this conformation appears to be shared by oxidation states in both the turnover and reactivation cycles, we propose that the resting state represents the second major conformation of MetH that was initially proposed 25 years ago^12^. Based on the fact that the cobalamin cofactor must interact with three distantly located active sites, it has been known that MetH must be a flexible enzyme^12,13,20,27,33^. Our work reveals that in the resting state, the majority of the enzyme’s flexibility is localized to the AdoMet domain. This model explains, for the first time, the tryptic proteolysis pathway that was shared by all oxidation states in the absence of flavoxin^12^, and why MetH can function as a three-domain enzyme^24,39^.

Interestingly, we found that CH_3_-H_4_folate is the only substrate that can induce a measurable conformational change from the resting state in both the Cob(II) and CH_3_-Cob(III) enzymes. Although in the case of the CH_3_-Cob(III) enzyme, X-ray exposure unavoidably led to reduction of the cofactor, this does not necessarily mean that the conformational change occurred in the Cob(II) state. Rather, it suggests that CH_3_-H_4_folate must have at least initiated a conformational change in the CH_3_-Cob(III) state to make homolysis of the Co-C bond irreversible. As was proposed in previous photolysis studies^23^ such a conformational change is likely to involve the uncapping of the B_12_ domain, which makes the methyl radical more susceptible to escaping the enzyme. By studying the 98 kDa fragment of *E. coli* MetH, we confirmed that the CH_3_-H_4_folate-induced conformation is indeed consistent with a cap-off B_12_ domain interacting with the folate domain. Furthermore, our cryo-EM model shows that in the resting state, the only active site that directly interacts with the B_12_ domain is that of the folate domain. Specifically, the cap subdomain and CH_3_-H_4_folate bind the same structural element in a mutually exclusive manner. CH_3_-H_4_folate is therefore the only substrate that can be rationalized to cause a conformational change from the observed resting state. Likewise, the existence of the resting-state conformation explains why MetH is able to sense CH_3_-H_4_folate. This provides a structural explanation for seminal findings from Banerjee, et al. that demonstrated that MetH prefers to carry out its two primary turnover reactions in a sequential order, with CH_3_-H_4_folate binding first and H_4_folate dissociating last^20^. Furthermore, sensitivity to CH_3_-H_4_folate provides MetH with an essential function in one-carbon metabolism. Although MetH serves as the link between the methionine and folate cycles, the enzyme is likely to be saturated with homocysteine in the cell^20^, and production of methionine is not an essential function as it can be obtained from diet^41^. In contrast, MetH is essential for assimilating CH_3_-H_4_folate into other folate species^11^, and thus, it would make sense for this substrate to have a role in conformationally gating enzyme turnover.

Given that oxidation states in both the turnover and reactivation cycles appear to share a common resting-state conformation, a key question is how the enzyme is able to switch between the two cycles. Previous spectroscopic studies^26^ indicated that flavodoxin must be able to bind Cob(II) MetH as it causes a change in the coordination geometry of the cobalamin cofactor from 5-coordinate (His-on) to 4-coordinate (His-off), while it is not able to bind the CH_3_-Cob(III) enzyme or reorganize its cofactor environment. NMR, electron transfer and cross-linking studies^12,22,26,42^ have further shown that the C-terminal AdoMet domain is required for the two proteins to interact and that a large conformational change must occur in the Cob(II) state to enable rapid reduction by flavodoxin. Later crystallographic studies established that a reactivation-state conformation of MetH exists^14,15,18^ and that this conformation is stabilized by the change in the coordination geometry of Cob(II) cofactor, which frees His759 to interact instead with residues in the AdoMet domain^18^. In this study, we have provided the first structural evidence that flavodoxin binding indeed favors the Cob(II) state over CH_3_-Cob(III). Under the condition of our study, which was near the pKa of free histidine, we found that both Cob(II) and CH_3_-Cob(III) MetH remain in a conformation that is largely consistent with the resting state at pH 7.6. However, a subtle change was observed in the Cob(II) scattering in lowering the pH from 7.6 to 6 that was accompanied by a shift in the UV-Vis absorption that was indicative of an increase in the His-off population. Based on these results, we hypothesize that although free MetH is predominantly in the resting-state conformation in both the Cob(II) and CH_3_-Cob(III) states, it more readily interconverts with the reactivation-state conformation in the Cob(II) state because the lower His759 ligand is able to interact with either the cofactor or the AdoMet domain. We further propose that flavodoxin is excluded in the resting-state conformation and favors interactions with the two C-terminal domains in the reactivation-state conformation. In this way, flavodoxin binding would be able to push the Cob(II) MetH population towards the His-off state.

Based on the findings from this study, we propose the following model for the mechanism by which MetH is able to switch between different functional modes. In this model, the resting-state conformation serves as the favored state in the absence of perturbations. During turnover, the enzyme will need to sample at least two additional conformations. In the resting state, the B_12_ Rossmann subdomain is already in contact with the back of the Hcy domain. It is plausible that the resting state of the enzyme is able to interconvert with a conformational state competent for homocysteine methylation without a major domain rearrangement. A possible trajectory for such a transition is populated by the collection of AlphaFold2 models in Figure S16A-B. In contrast, our results indicate that resting state cannot easily transition to folate-demethylation state without CH_3_-H_4_folate binding. However, when CH_3_-H_4_folate binds to the folate domain, it engages the folate-sensing loop in a way that undocks the bound cap subdomain and prevents the resting state from reforming. Such a mechanism may serve to rebalance the conformational distribution of the enzyme such that both turnover states are sampled. Finally, as described above, we propose that the resting state can interconvert with the reactivation state and that this forward conversion is more favored when His759 can participate in inter-domain interactions. Thus, although Cob(II) MetH favors the resting state conformation, it can be recognized by flavodoxin when it occasionally samples the His-off, reactivation conformation. In this way, flavodoxin can syphon the inactive enzyme from the turnover-cycle ensemble to initiate reactivation, as shown in Figure 1B. Once CH_3_-Cob(III) is regenerated by reductive methylation, the enzyme would again favor the resting-state conformation. By having a common resting-state conformation, the enzyme would be able to “reset” and return to the turnover cycle.

Over the course of this study, several technical advances emerged that enabled key aspects of our investigation. These included the rapid evolution of single-particle cryo-EM^43^ as well as the development of anoxic SAXS. On the computational front, the advent of AlphaFold2^32^ had a surprising impact. We showed that AlphaFold2 succeeded in predicting the resting state, a conformation that does not yet exist in the training dataset. Although structure prediction cannot replace experimental structural biology, this was an extraordinary feat. We further demonstrated two unique uses of the AlphaFold2 database that greatly expand upon conventional analyses of multiple sequence alignments (MSAs) in structural biology. First, by analyzing the collection of predicted models in the public AlphaFold2 database, we were able to visualize conformations that differ significantly from those represented in the PDB (Figure S16). In the case of MetH, sequences with high similarity to the *E. coli* enzyme are more likely to be predicted in the reactivation state, which is highly represented in the PDB (Figure S20A). Thus, the availability of structure predictions for a diverse sequence dataset is likely to reveal conformations that have never been considered before (Figure S20B-E), especially for flexible, muti-domain enzymes. Second, analysis of the AlphaFold2 database also provided insight into how sequence variability can still lead to conserved functions (Figure S14). Based on conventional MSA analysis, we found that residues involved in the interface between the cap subdomain and the folate domain are not fully conserved. Analysis of alternate sequence motifs in the AlphaFold2 database, however, suggested that the non-conserved residues have co-evolved in a way that conserves the overall set of interactions. Such an insight cannot be easily obtained from sequence analyses alone. It is worth repeating that structural biology should not be replaced by structure prediction, which does not yet have angstrom to sub-angstrom level accuracy. However, we anticipate that structure prediction will continue to enhance experimental efforts, possibly in ways that we cannot yet imagine.

## Materials and Methods

Untagged, full-length *E. coli* MetH (residues 2-1227) and its fragment containing the three N-terminal domains (residues 2-896) were prepared following previously described methods^12,31,44^. His6-tagged *E. coli* flavodoxin was purified by a cobalt-affinity column followed by size exclusion. Tagless *T. filiformis* MetH was prepared using a cleavable His6-SUMO tag and reconstituted as described in detail in the SI Appendix. Detailed descriptions of protein expression and purification, preparation of pure oxidation states, cofactor reconstitution, limited proteolysis, SAXS, cryo-EM, bioinformatics, and AlphaFold2 analyses are available in the SI Appendix. The oxidation state and coordination geometry of the cobalamin cofactor was verified by UV-Vis spectroscopy for every sample in every experiment. Unless otherwise specified (e.g., for experiments performed at pH 6), CH_3_-Cob(III) and Cob(II) states were verified to be almost entirely His-on for both the *E. coli* and *T. filiformis* enzymes.

## Data Availability

The cryo-EM map has been deposited in the Electron Microscopy Data Bank under accession code EMD-8G3H and the model has been deposited in the Protein Data Bank under accession code EMD-29699.

## Acknowledgements and Funding Sources

The authors are grateful to Prof. Rebecca Taurog (Williams College) for providing us with plasmids, reagents, and helpful tips; to Gabrielle Illava for setting up the anoxic SAXS system; and to Dr. Amanda Byer for assistance with SAXS data collection and critical reading of this manuscript. We are also grateful to Drs. Richard Gillilan, Qingqiu Huang and Jesse Hopkins for assistance with SAXS setup at CHESS, and Drs. Katie Spoth and Mariena Ramos (CCMR) as well as Drs. Edward Eng and Eugene Chua (NCCAT) for assistance with cryo-EM data collection. Finally, we thank Prof. Yan Kung (Bryn Mawr) for sharing a protocol for titanium(III) citrate, Prof. Dominica Borek (UT Southwestern) for suggesting the use of apoferritin, and Prof. Markos Koutmos (U Mich) for suggesting the use of thermophilic enzymes. SAXS was performed at CHESS, which is supported by award P30 GM12416601 (to MacCHESS) from the National Institute of General Medical Sciences, National Institutes of Health (NIH) and by the NY State Empire State Development Corporation (NYSTAR). Additional SAXS was performed with a Xenocs BioXolver funded by NIH grant S10OD028617. Cryo-EM data collection was performed at the Cornell Center for Materials Research (CCMR), which is supported by the National Science Foundation (NSF) under award DMR-1829070, as well as at the National Center for Cryo-EM Access and Training (NCCAT) in New York City, which is supported by the NIH Common Fund Transformative High Resolution Cryo-Electron Microscopy program grant number U24 GM129539. This work was supported by NIH grants GM100008 and GM124847 and startup funds from Princeton University and Cornell University to N.A.

## Author Contributions

N.A. conceived of the project, oversaw and supervised the research, analyzed data, and acquired funding. M.B.W. designed research and performed the majority of experiments and data analysis. H.W. prepared the 98 kDa fragment, performed SAXS on the fragment, measured UV-Vis spectra, and performed AlphaFold2 analysis. A.A.B. performed bioinformatics. M.B.W. and N.A. wrote the manuscript with contributions from H.W. and A.A.B. All authors edited the manuscript.

