## Supporting Information for "Conformational switching and flexibility in cobalamin-dependent methionine synthase studied by small-angle X-ray scattering and cryo-electron microscopy"

**This PDF file includes:**

- Extended Methods
- Tables S1-S11
- Figures S1-S20
- Supporting References

**Other supporting materials for this manuscript include the following:**

- Movies S1

### Extended Methods

**Expression and purification of *E. coli* MetH:** *E. coli* XL1-Blue cells transformed with the pMMA-07 plasmid encoding for untagged, full-length *E. coli* MetH (residues 2-1227)[1] were kindly provided by Prof. Rebecca Taurog (Williams College). The enzyme was expressed from resulting glycerol stocks in cobalamin-containing minimal media and purified following previously described methods[2,3], with some modifications. Reagents were purchased from Sigma or ThermoFisher unless otherwise specified.

For expression, we adapted a previously described protocol that uses ethanolamine as a carbon and nitrogen source as a way to enhance intracellular cobalamin concentrations[3]. A minimal growth medium (buffered at pH 7.4 and containing all 20 amino acids, ethanolamine, hydroxocobalamin, zinc, and other essential nutrients) was prepared in the same manner as previously described[3] and supplemented with 100 µg/mL ampicillin. A 500-mL starter culture was grown in the growth medium overnight at 37 °C, 250 rpm from a glycerol stock prepared from a single colony. Large-scale growth media were inoculated with starter culture (25 mL/1 L), and the cultures were grown at 37 °C, 250 rpm to an $OD_{600}$ of between 0.5 and 0.6. Expression was induced with 1 mM isopropyl β-d-1-thiogalactopyranoside (IPTG) for ~18 hours at 18 °C, 250 rpm. Cells were harvested by centrifugation at 4500 × g for 15 minutes at 4 °C.

The lysis buffer was 10 mM potassium phosphate (KPi) pH 7.2, 1 mM tris(2-carboxyethyl)phosphine (TCEP). Cells were resuspended in lysis buffer supplemented with cOmplete mini protease inhibitor cocktail (Roche, 1 pellet/10 mL) and DNAse (Alfa Aesar, 17 units/10 mL) and lysed by passage through an Avestin Emulsiflex C3 homogenizer at 4 °C for 10-15 minutes. Resultant lysate was centrifuged at 33,000 × g for 40 minutes at 4 °C, and the supernatant was passed through a 0.4-µm filter and loaded onto a HiPrep DEAE anion exchange column. The column was washed with 10 column volumes (CVs) of lysis buffer, and bound protein was eluted with a 20-CV linear gradient from 10 to 500 mM KPi, pH 7.2, 1 mM TCEP. Protein-containing fractions were exchanged into lysis buffer using a HiPrep 26/10 desalting column and loaded onto a Mono-Q HR 16/10 anion-exchange column. Protein was eluted with a 20-CV linear gradient from 100 to 320 mM KPi pH 7.2, 1 mM TCEP. Following elution, the protein was loaded onto a HiLoad 16/600 Superdex 200-PG size-exclusion column pre-equilibrated with the final buffer (50 mM HEPES pH 7.6, 150 mM NaCl, with 1-2.5 mM dithiothreitol (DTT)). The purest non-aggregate fractions (as assessed by SDS-PAGE) were pooled, concentrated and frozen in liquid $N_2$, and stored at –80 °C. The entire purification was performed at 4 °C. For all buffers except the final one, TCEP was added the morning of use, and DTT was added to the final buffer immediately before use. As the purification protocol takes two full days, protein was stored overnight at 4 °C after the HiPrep 16/10 desalting step to avoid unnecessary freeze-thawing.

**Preparation of pure oxidation states of *E. coli* MetH:** MetH purifies as a mixture of states (primarily methylcobalamin or $CH_3$-Cob(III), Cob(II) and the non-physiological hydroxocobalamin OH-Cob(III))[2]. However, the mixture can be reductively methylated to the methylcobalamin form, from which the enzyme can be prepared in each of the other physiological oxidation states and identified by their unique UV-Vis absorption spectra. Purified oxidation states were prepared in a positive-pressure $N_2$-atmosphere mBraun anaerobic chamber ($O_2$ < 0.1 ppm, 15 °C) in a darkened room, with the only source of illumination being a red lamp designed for photographic film development to prevent unwanted methylcobalamin photolysis.

To prepare titanium (III) citrate solution, a saturated solution of sodium bicarbonate and a solution of 1 M Tris pH 7.0 were prepared and deoxygenated on a Schlenk line. Solutions were brought into the mBraun chamber along with deoxygenated deionized water and unopened commercial titanium chloride ($TiCl_3$, 20% solution in 2 N HCl) from EMD Millipore. 5.8 g sodium citrate was weighed, brought into the chamber, and dissolved in 20 mL of water while stirring. 7.5 mL of titanium chloride solution was added, and the pH was adjusted approximately to 7.0 using sodium bicarbonate. The concentration of the resulting titanium(III) citrate solution was then was quantified with methylviologen. A 100-µL aliquot was prepared by diluting the titanium(III) citrate solution 10-fold in Tris buffer, and 1 µL of the diluted solution was mixed with 500 µL of 10 mM methylviologen, which reduced and turned blue. The concentration of the titanium (III) citrate was then related to the extinction coefficient of reduced methyl viologen (9.59 $mM^{-1}$ $cm^{-1}$ at 578 nm)[4]. The remaining, undiluted titanium (III) citrate solution was then adjusted to a final concentration of 50 mM in 100 mM Tris pH 7.0.

Methylcobalamin enzyme was prepared from the as-isolated MetH, similarly as described previously[2]. As-isolated MetH was brought frozen into the mBraun chamber, then thawed and equilibrated with the atmosphere on ice or a cold block for at least 1 hour under anoxic conditions. Per 100 nmol of protein (as estimated by a sequence-based theoretical extinction coefficient of 136,600 $M^{-1}$ $cm^{-1}$ at 280 nm), 15 µL of *S*-adenosylmethionine (38 mM in 1 mM HCl) was slowly added to the protein by drop-pipetting on the side of the tube. Following addition, 50 µL of titanium(III) citrate solution (50 mM in 100 mM Tris HCl, pH 7.0) was slowly added to a 5- or 10-fold excess of the protein concentration. The reaction was monitored via UV-Vis absorption using an Implen Nanophotometer or ThermoFisher Nanodrop in microvolume mode and typically took 5-10 minutes. The disappearance of the shouldered absorption peaks at ~477 nm, and the strengthening of the broad peak across 528 nm were indicative of conversion to methylcobalamin. The resulting methylcobalamin enzyme was exchanged into final buffer using either GE/Cytivia PD-10 desalting columns or Bio-Rad MiroSpin columns (depending on the volume of protein solution) and concentrated.

Following buffer exchange, the protein concentration determined by the theoretical extinction coefficient at 280 nm typically correlated to within ~10% of the concentration of the cofactor as estimated by the methylcobalamin extinction coefficient (8910 $M^{-1}$ $cm^{-1}$ at 525 nm)[5], and the theoretical extinction coefficient at 280 nm was used for protein quantification in further experiments.

The turnover-cycle Cob(I) enzyme was prepared by anaerobic homocysteine (Hcy) demethylation of methylcobalamin enzyme, which was performed inside the mBraun chamber. A 5-fold molar excess of anaerobic Hcy solution (10 or 100 mM in MetH buffer: 50 mM HEPES pH 7.6, 150 mM NaCl, 2.5 mM DTT) was added to the methylcobalamin enzyme, which was typically at ~10-40 µM. The formation of the Cob(I) species was monitored by the appearance of the absorption peak at 390 nm and the disappearance of the broad peak at 528 nm and the reaction typically proceeded to completion rapidly. Cob(I) enzyme was then exchanged back into MetH buffer and concentrated if necessary.

Cob(II) enzyme was prepared by aerobic Hcy demethylation of methylcobalamin enzyme. DTT was added to methylcobalamin enzyme to a final concentration of 10 mM inside the mBraun chamber. A 5-fold molar excess of anaerobic Hcy solution (10 or 100 mM in MetH buffer) was then added to the reaction mixture, and the solution was mixed and allowed to react for ~5 minutes and monitored for demethylation via UV-Vis absorption. The mixture was then briefly brought out of the glove box and exposed to ambient atmosphere for 2-3 seconds, closed again and then inverted several times. The solution was then brought back into the glove box, and oxidation to Cob(II) was verified by the disappearance of the Cob(I) peak at 390 nm and appearance of the Cob(II) peak at ~477 nm. Cob(II) enzyme was then exchanged into MetH buffer and concentrated if necessary.

Cob(I) enzyme was not stable in solution for long periods of time (> 1-2 hours) even under rigorously anoxic conditions, and those not prepared immediately prior to experiments typically oxidized and degraded. On the other hand, the oxidation states of the Cob(II) and methylcobalamin enzymes were reasonably stable in buffer containing 1-2.5 mM DTT for many hours at 4 °C, although the methylcobalamin enzyme slowly photolyzes in ambient light over time and as such needed to be protected from light. To enable accurate background subtraction of SAXS data, enzyme aliquots were frozen concurrently with the exact buffer used for the final buffer exchange step. Cob(I) enzyme for SAXS experiments was prepared in an on-site Coy anaerobic chamber immediately before measurement and never freeze-thawed. The successful conversion of the as-isolated *E. coli* MetH from one oxidation state to another (as assessed by UV-Vis absorption spectroscopy) was used as an indication of active enzyme[2,6].

**Expression and purification of *E. coli* flavodoxin FldA:** A pET28 vector encoding for C-terminal hexa-histidine ($His_6$)-tagged *E. coli* flavodoxin (FldA) was kindly provided by Prof. Rebecca Taurog (Williams College). A glycerol stock was prepared from a single colony of *E. coli* BL21 (DE3) cells transformed with the plasmid. A 25-mL starter culture in LB (Miller) media containing 50 µg/uL kanamycin was grown from the glycerol stock overnight at 37 °C, 250 rpm. 1 L of LB (Miller) media containing 50 µg/mL kanamycin was inoculated with 5 mL of starter culture, and the culture was grown to an $OD_{600}$ of 0.6 at 37 °C, 250 rpm. Expression was then induced with 1 mM IPTG for 4 hours at 30 °C, 250 rpm. Cells were harvested at 4500 × g for 15 minutes at 4 °C and re-suspended in lysis buffer (20 mM sodium phosphate (NaPi), 40 mM imidazole pH 7.4) supplemented with cOmplete mini Protease inhibitor cocktail (1 pellet/10 mL) and DNAse (Alfa Aesar, 17 units/10 mL). The cells were lysed by passage through an Avestin Emulsiflex C3 homogenizer at 4 °C for 10-15 minutes. Lysate was centrifuged for 40 minutes at 33,000 × g at 4 °C, and the supernatant was passed through a 0.4-µm filter and loaded onto a Ni-affinity column (HisTrap FF crude, 5 mL). The column was then washed with 10 CVs of lysis buffer, and the protein was eluted with a 20-CV linear gradient from 40 to 300 mM imidazole (in 20 mM KPi, pH 7.4). Following elution, the protein-containing fractions were combined and loaded onto a HiLoad 16/600 Superdex 75-PG size-exclusion column pre-equilibrated in final buffer (20 mM KPi, pH 7.4). Eluting fractions were checked for purity by SDS-PAGE, and the purest non-aggregate fractions were pooled, concentrated and frozen in liquid $N_2$. *E. coli* FldA purifies with endogenous flavin mononucleotide (FMN) bound. The final FldA concentration was checked in two ways: first, with the literature value[7,8] for oxidized FMN absorbance at 466 nm of 8,250 $M^{-1}$ $cm^{-1}$ and second, with the sequence-based theoretical protein extinction coefficient at 280 nm of 31,200 $M^{-1}$ $cm^{-1}$. The results agreed to within ~10%, and the theoretical extinction coefficient at 280 nm was used for protein quantification in further experiments.

**Expression and purification of *T. filiformis* MetH:** A pET-28c+ vector encoding for N-terminal $His_6$-SUMO-tagged *T. filiformis* MetH was synthesized by GenScript and transformed into *E. coli* T7 Express cells (NEB). A 10-mL starter culture was grown overnight at 37 °C, 250 rpm in LB (Miller) media from a glycerol stock prepared from a single colony. A cobalamin supplement solution was prepared in the same way as previously described for the *E. coli* enzyme[3]. For large-scale growth, the LB medium was supplemented with cobalamin supplement (13.5 mL/1 L) and inoculated with the starter culture (10 mL/1 L). The large-scale cultures were grown at 37 °C, 250 rpm to an $OD_{600}$ of between 0.5 and 0.6. Expression was induced with 1 mM IPTG for 4 hours at 37 °C, 250 rpm. Cells were harvested by centrifugation at 4500 × g for 15 minutes at 4 °C.

Cell paste was resuspended in lysis buffer (20 mM KPi pH 7.4, 1 mM TCEP) supplemented with cOmplete mini protease inihibitor cocktail (1 pellet/10 mL) and Benzoase (17 units/10 mL). The resuspended cells were lysed by passage through an Avestin Emulsiflex C3 homogenizer for 10-15 minutes at 4 °C. Resultant lysate was collected and centrifuged at 33,000 × g for 40 minutes at 4 °C, and the supernatant was passed through a 0.4-µm filter and loaded onto a cobalt affinity column (HiPrep TALON crude, 5 mL). The column was washed with 5 CVs of lysis buffer, and bound protein was eluted with a 10-CV linear gradient from 0 to 500 mM imidazole (in 20 mM KPi pH 7.4, 1 mM TCEP). Protein-containing fractions were exchanged back into lysis buffer using Cytivia/GE PD10 desalting columns. Pooled fractions were then incubated with a $His_6$-tagged SUMO protease (1:300 protease:MetH molar ratio) at room temperature for 2 hours. The mixture was then loaded onto a HiPrep TALON crude column. Cleaved, untagged protein was collected in the column flowthrough, and $His_6$-SUMO-containing protein was retained by the column. The untagged protein was concentrated and loaded onto a HiLoad 16/600 Superdex 200-PG size-exclusion column pre-equilibrated with final buffer (50 mM HEPES, 150 mM NaCl, 2.5 mM DTT pH 7.6). The purest non-aggregate fractions (as estimated by elution volume and SDS-PAGE) were pooled, concentrated and frozen in liquid $N_2$ and stored at –80° C. TCEP was added to buffers the morning of use, while DTT was added to the final buffer immediately before use. Cleavage of the $His_6$-SUMO tag yields the native protein sequence.

**Reconstitution of *T. filiformis* MetH:** *T. filiformis* Cob(II) enzyme was reconstituted from the as-isolated apoenzyme following methods previously reported for *Thermotoga maritima* MetH[9]. As-isolated apoenzyme solution was heated to 60 °C on a heat block and held at temperature for 2 minutes. A 10-fold molar excess of hydroxocobalamin (stock solution: 330 mM in 0.9 M NaCl) was then added and mixed, and the mixture was incubated for 3 minutes at 60 °C. Following this, the mixture was then exchanged back into the final purification buffer for immediate experimental use. Reconstitution with hydroxocobalamin in DTT-containing buffer resulted in reduction to Cob(II) enzyme after ~5 minutes, as monitored by UV-Vis absorption. Reconstitution appeared to be 1:1, as estimated by comparison of the sequence-based theoretical extinction coefficient of the protein at 280 nm of 123,080 $M^{-1}$ $cm^{-1}$ to the calculated extinction coefficient of free cob(II)alamin (5030 $M^{-1}$ $cm^{-1}$ at 472 nm). The Cob(II)alamin extinction coefficient was calculated by preparing a reference solution of free hydroxocobalamin, which was then reduced to Cob(II)alamin by addition of DTT to a final concentration 10 mM.

**Purification of 3-domain Cob(II) *E.coli* MetH:** A 3-domain fragment of Cob(II) *E.coli* MetH was generated via proteolysis of full-length Cob(II) *E.coli* MetH. 800 µL of 24 µM full-length MetH was incubated with trypsin (final concentration 0.005 mg/mL) at room temperature for 32 min in 50 mM potassium phosphate buffer, pH 7.2, 1 mM TCEP. The proteolysis was quenched by addition of phenylmethylsulfonyl fluoride (PMSF) to a final concentration 0.25 mM, and the reaction tube was subsequently placed on ice. The proteolysis yielded primarily the 98 kDa and 38 kDa fragments as previously observed[10], which correspond to the three N-terminal domains (residues 2-896) and the C-terminal AdoMet domain (residues 897-1227), respectively. An SDS-PAGE band at ~70-kDa was also observed, which is potentially a mixture of the N- and C-terminal halves, given the lack of an observed band for the $B_{12}$ domain alone (28 kDa). The mixture was centrifuged at 21,130 × g for 6 min at 4 °C and then loaded onto a Mono Q 5/50 GL anion-exchange column, and the fragments were eluted with a 20-CV linear gradient from 50 to 320 mM KPi pH 7.2, 1 mM TCEP at a flow rate of 0.5 mL/min. The 38 kDa fragment eluted first at ~ 100 mM KPi, and a mixture of the 98 kDa and 70 kDa fragments eluted at ~180 mM KPi. The latter mixture was left at 4 °C overnight, concentrated, and loaded onto a Superdex 200 10/300 GL column pre-equilibrated with final MetH purification buffer (50 mM HEPES, 150 mM NaCl, 2.5 mM DTT pH 7.6) the next day. The purest non-aggregate fraction without contamination of the 70 kDa fragment(s) (as assessed by SDS-PAGE, Figure S17B) was concentrated to 25 µM (overall yield of 13%), frozen in liquid $N_2$ and stored at –80 °C.

**Size exclusion chromatography-coupled SAXS (SEC-SAXS) of *E. coli* MetH:** SEC-SAXS experiments were performed at the ID7A1 station of the Cornell High Energy Synchrotron Source (CHESS). Samples were centrifuged for 10 minutes at 14,000 × g (4 °C) and loaded onto a Superdex 200 5/150 GL (GE/Cytiva) column, operated by an AKTA Pure FPLC (GE) at 4 °C. The sample was eluted at a flow rate of 0.2 mL/min directly into the *in vacuo* X-ray flow cell (also at 4 °C) after it passed through the UV monitor. X-ray scattering was recorded on a Pilatus 100K (Dectris) detector using a ~9.9 keV 250 × 250 µm X-ray beam, with an attenuated flux of ~2 × $10^{11}$ photons/sec. The detector was placed 1.53 m from the sample and recorded data over a $q$-range of 0.0091 to 0.275 $Å^{-1}$ with the momentum transfer vector defined as $q = (4\pi/\lambda)\sin(\theta)$, where $\lambda$ is the X-ray wavelength and $2\theta$ is the scattering angle. Roughly 500 to 600 2-second exposures were acquired over the entire elution (Table S1). A buffer scattering profile was created by averaging scattering profiles preceding the elution peak and was subtracted from exposures selected from the elution peak to create the $I(q)$ vs $q$ curves used for subsequent analyses. Data were reduced and processed initially (including deconvolution by evolving factor analysis[11]) in BioXTAS RAW[12] and then further analyzed using custom scripts written in MATLAB and Python.

**Batch-mode SAXS of *E coli* MetH:** Batch-mode experiments on the *E. coli* MetH were performed at the ID7A1 station of the Cornell High Energy Synchrotron Source (CHESS) with sample oscillation in an *in vacuo* X-ray cell. Images were recorded on an Eiger 4M detector using a ~9-14 keV 250 × 250 µm X-ray beam with a flux of ~6-15 x $10^{11}$ photons/sec. The sample-detector distance was ~1.7 m, and the detector collected data over a *q*-range of ~0.008 to ~0.5 $Å^{-1}$. Data were reduced and processed initially in BioXTAS RAW[12] and then further analyzed using custom scripts written in MATLAB and Python.

Batch-mode samples of the full-length enzyme were prepared at 20 µM concentration and were centrifuged at 10,000 × g for 10 minutes at 4 °C immediately before loading 30-40 µL into the X-ray cell set at 20 °C. For experiments with methylcobalamin enzyme, the X-ray hutch was fully darkened and protected from outside light with a laser curtain, and the sample cell was illuminated only with the same red lamp used for methylcobalamin enzyme preparation. For experiments with Cob(I) enzyme, the entire SAXS experimental setup was contained in a fully anaerobic loop, where the tubing from the loading channel to the sample cell and again from the sample cell to the waste were terminated within an anaerobic chamber (Coy laboratories, 95%/5% $N_2/H_2$ atmosphere, $O_2$ < 30 ppm). The tubing was flushed with 100 mM sodium dithionite prior to loading experimental samples in order to scavenge any residual oxygen. Cob(I) enzyme was prepared fresh from methylcobalamin enzyme immediately prior to X-ray exposure in the same chamber, following the protocol described above. The integrity of the oxidation state was assessed by UV-Vis absorption measurements performed inside the anaerobic chamber using an Implen Nanophotometer in nanovolume mode. Specific data collection details for the purified oxidation states can be found in Table S2. Where possible, reported dissociation constants ($K_D$ = 1.2 µM[10] for AdoMet, $K_D$ = 20.8 µM[13] for AdoHcy, $K_D$ = 10-20 µM for Hcy)[14] were used to achieve saturation of substrate/product binding.

For the folate titration experiment, enzymatically prepared[15] enantiomerically pure (6*S*)-5-$CH_3H_4$-PteGlu$_3$ solution (3.4-13.4 mM in 10 mM BME) was obtained from Prof. Becky Taurog (Willams College). The same volume of (6*S*)-5-$CH_3H_4$-PteGlu$_3$ solution was added to both MetH solution and the buffer blank and incubated for 2 minutes at room temperature prior to centrifugation (Table S3). For the flavodoxin titrations, MetH and FldA were buffer-exchanged at the beamline into 44 mM sodium/potassium phosphate pH 6.0, 1 mM DTT (ionic strength normalized to 0.15) prior to mixing and were incubated together for 2 minutes at room temperature prior to centrifugation (Tables S4-5). In Figure 3, FldA is abbreviated as "Fld".

For experiments with the 3-domain *E. coli* proteolytic fragment, samples were centrifuged at 21,130 × g for 10 minutes at 4 °C immediately prior to loading into the X-ray cell set at 4 °C. Control experiments showed that folate-dependent behavior was unchanged at the lower temperature for the full-length enzyme. The (6*R*,*S*)-$CH_3H_4$-PteGlu$_3$ substrate used for these experiments was purchased from Schrick's Labs, and 12.1 mM solutions in 10 mM BME were prepared in an anaerobic chamber (Coy laboratories, $O_2$ < 30 ppm). Specific data collection and analysis details for this experiment can be found in Table S10.

**Batch-mode SAXS of *T. filiformis* MetH:** SAXS data collection for *T. filiformis* MetH was performed at a wavelength of 1.542 Å on a Xenocs BioXolver instrument equipped with a rotating anode X-ray source and a Pilatus 300K detector. Protein at a concentration of 20 µM was loaded via autoloader (from a 96-well plate kept at 4 °C) into the *in vacuo* sample cell in batch mode at 20 °C, and data was collected over a *q*-range of 0.0005-0.439 $Å^{-1}$. The low-*q* data close to the transmitted beam is not usable, and thus the *q*-range used for analyses was 0.0146-0.439 $Å^{-1}$. 10 individual frames (120 seconds each) were averaged per buffer or protein sample. Samples incubated with and without (6*S*)-5-$CH_3H_4$-$PteGlu_3$ were prepared identically to the corresponding full-length *E. coli* MetH samples, as described above. Data were initially processed in BioXTAS RAW[12] (Table S6).

**SAXS data processing and analysis:** SAXS data were processed initially in BioXTAS RAW[12] and then further analyzed using custom scripts written in MATLAB and Python. The pair-distance distribution, $P(r)$, was calculated from the indirect Fourier transform of the scattering intensity $I(q)$ in GNOM[16]. The ensemble optimization method (EOM)[17,18] was used to build ensembles and assess the flexibility of the full-length MetH in solution. An *E. coli* resting-state model for the 3 N-terminal domains (residues 3-896) was built by aligning the AlphaFold2[19] model for the two N-terminal domains and the crystal structure of the cap-on $B_{12}$ domain (PDB 1BMT[20]) to our *T. filiformis* EM structure. This model and a crystal structure of the C-terminal AdoMet domain (PDB 1MSK[21] with ligands removed; residues 901-1227) were treated as two flexibly linked rigid bodies. A pool of 10,000 structures sampling the conformational space was generated using RANCH[17,18], and subsets were fit to experimental SAXS data obtained on the full-length *E. coli* MetH using the program GAJOE[17,18]. Fit statistics are shown in Table S9. All displayed SAXS profiles (and fits) were normalized by the experimental Guinier $I(0)$ value; Kratky plots were additionally scaled by a factor of $10^4$ for visual clarity.

**Sequence analysis:** The EFI-EST server[22] was used to generate a sequence-similarity network (SSN) from 8,047 MetH sequences using Pfam[23] accession PF02965 and to perform further analysis (including coloring and sequence conservation). The network was visualized in Cytoscape[24], with the edge alignment score was set to 450 corresponding to a minimum identity of 49%. The 8,047 MetH sequences were manually curated for outlier and redundant sequences above 90% threshold to 5,520 sequences and aligned with MAFFT[25] using the FFT-NS-1 option. Of the 5,520 sequences, 1295 have a Phe residue at position 720 and a Gln at position 717 (*T. filiformis* numbering), while 3501 have a Gln at position 720 and a Phe at position 717. *T. filiformis* MetH is in the former group, while *E. coli* MetH is in the latter. Sequence logos were calculated for these two groups in Skylign[26] using the default options.

**Cryo-EM sample preparation and data collection:** *T. filiformis* Cob(II) MetH was mixed with horse spleen apoferritin, both in 50 mM HEPES, 150 mM NaCl, 2.5 mM DTT pH 7.6, at a final concentration of 2 µM for each protein. 4 µL of the protein mixture was applied to glow-discharged Quantifoil R1.2/1.3 holey carbon support grids, which were blotted on both sides and frozen in liquid ethane using a ThermoFisher Vitrobot™ Mk. IV (chamber set to 4° C and 100% humidity, blot time of 3.5 seconds, blot force of 3, and wait time of 0). Grids were screened on a 200 keV Talos Arctica operated by the Cornell Center for Materials Research (CCMR), and a full dataset was collected at the National Center for CryoEM Access and Training (NCCAT). Images were collected over two identically prepared grids on a 300 keV Titan Krios equipped with a Gatan Biocontinuum Imaging energy filter (set to 30 eV width) and a 24-megapixel Gatan K3 detector. In total, 12,768 50-frame movies were collected at a nominal pixel size of 1.07 Å and at a dose rate of 61.7 e- /Å$^2$. Additional data collection details are described in Table S7.

**Cryo-EM data processing:** An initial dataset containing 12,768 micrographs was manually curated to a final set of 9,656 micrographs. Patch motion correction and patch CTF estimation were performed on the curated micrograph stack using the default implementations in cryoSPARC V3.2[27]. Particles were picked in cryoSPARC using templates generated from 2D classification of ~500 manually picked particles, yielding an initial particle stack of 9,994,870 particles. The majority of these (~70%) represented apoferritin particle picks and were manually removed after initial reference-free 2D classification. The remaining particles were subjected to *ab initio* reconstruction and heterogeneous refinement in cryoSPARC using 3 classes, which separated the particle stack into a class resembling the species of interest, a class containing unknown heterogeneous/blurry contrast particles, and a class representing just the two N-terminal domains. Particles containing the species of interest were further filtered by reference-free 2D classification followed by another round of *ab-initio* reconstruction and heterogeneous refinement with 2 classes, which cleanly separated the particle stack into one class appearing to contain only the two N-terminal domains and another containing the species of interest. Full-size particles were extracted at a box size of 384 pixels (~411 Å) prior to the second round of *ab-initio* reconstruction and heterogeneous refinement. Homogenous refinement of the final class of interest, followed by removal of potential duplicate particle picks and non-uniform refinement (without per-particle defocus or per-group CTF, as both were found to hurt resolution or map quality) in cryoSPARC yielded a 3.60-Å map (by FSC=0.143) from 257,706 particles. Data collection details can be found in the Table S7, and the processing pipeline is additionally visualized in Figure S9.

**Cryo-EM model building and refinement:** An initial starting model was built by rigid-body fitting in Chimera[28] using models for the individual domains taken from the AlphaFold2[19] rank-1 model for the *T. filiformis* MetH sequence. The resulting model was subjected to real-space rigid-body refinement in PHENIX[29] and then migrated to ChimeraX[30] for ISOLDE[31] modeling. The cobalamin ligand was not modeled at this stage as ISOLDE does not currently contain parameterization for it. A global simulation was allowed to run and settle on ISOLDE. Following concurrent manual model rebuilding and local simulations, another round of global simulation was performed to alleviate remaining clashes. The cobalamin ligand was docked in by alignment of the 1BMT[20] structure to the His759-containing cobalamin-binding region of the $B_{12}$-binding subdomain, and the methyl group was removed for consistency with the experimental oxidation state. The $B_{12}$ position was then refined by real-space rigid-body refinement in PHENIX. Several subsequent rounds of manual rebuilding in COOT were required; in particular, substantial rebuilding of the β-sheets surrounding the dimethylbenzimidazole was necessary to alleviate clashes introduced by addition of the cobalamin ligand, as it was not present during ISOLDE simulation. Map and model statistics can be found in Tables S7 and S8, respectively.

**Generation of 3-domain MetH models for fitting SAXS data:** Because all AlphaFold2[19] predictions of *E. coli* MetH depict the reactivation conformation, homology models were constructed for the resting state, homocysteine state, and folate state. Residues 9-896 of the *E. coli* MetH sequence were used for homology modeling based on proteolysis and mass spectrometry data[10]. The resting-state homology model was made with SWISSmodel[32] using the *T. filiformis* cryo-EM structure as the template. Likewise, the homocysteine-state and folate-state homology models were made using publicly available *Mus musculus* (UniProt A6H5Y3) and *Sphaerochaeta sp.* (UniProt A0A521IQ66) AlphaFold2[19] models, respectively, as templates. In all homology models, a methylcobalamin ligand was placed in the models by alignment of the 1BMT[33] structure to the His759-containing cobalamin-binding loop of each model. To match the experimental Cob(II) oxidation state, the methyl group was then removed from the cofactor. The missing N-terminal residues (2-8) were modeled while fitting to SAXS data on the 3-domain fragment using the program CORAL[34], and the resulting theoretical scattering profile and fit were re-calculated profile using CRYSOL (99 spherical harmonics, constant subtraction allowed). Fit statistics can be found in Table S11.

**AlphaFold2 structure prediction:** AlphaFold2[19] predictions were performed on a local installation using the default five model parameters and a max template date set to 2020-05-14. For the *T. filiformis* MetH enzyme, the sequence from UniProt accession number A0A0A2XCD7 was used, consistent with the experimental construct. Prediction was run on the full-length and truncated (residues 1-878) sequences. For the

*T. maritima* MetH enzyme, the sequence from UniProt accession TM0268 was used. Additional models were obtained from the public AlphaFold2[19] database as described below.

**Analysis and clustering of AlphaFold2 models:** To analyze MetH conformations predicted by AlphaFold2[19], sequences between 1100 and 1300 amino acids in length (to filter out most incomplete or 3-domain predictions) were selected from the MetH SSN (Figure S7). Sequences not containing full domain annotations and full binding-site annotations for zinc, cobalamin and AdoMet from UniProt were removed. AlphaFold2[19] predictions of the curated 4,915 MetH sequences were then downloaded from the public Alphafold2 database and aligned by the two N-terminal domains to the *E. coli* prediction result (UniProt P13009) using custom Python scripts and PyMol tools. Python scripts were written to perform conformational clustering of the models using the PyMol align command with outlier rejections enabled. To count the number of models in the resting-state conformation, models with Cα RMSD < 2.5 Å to the cap-on $B_{12}$ structure (PDB 1BMT) were kept and those with Cα RMSD < 3 Å to the reactivation-state structure (PDB 3BUL) were excluded. The result was then confirmed by the Cα RMSD distribution to the resting-state cryo-EM structure. To count the number of models in the homocysteine-state conformation, models with Cα RMSD < 2.5 Å to the 3 N-terminal domains of the *Mus musculus* prediction (UniProt A6H5Y3) were kept and those with Cα RMSD < 2.5 Å to the resting-state cryo-EM structure, Cα RMSD < 3 Å to the reactivation-state structure, or Cα RMSD < 3 Å to the cap-on $B_{12}$ structure were excluded. To count the number of models in the folate-state conformation, models with Cα RMSD < 2.5 Å to the 3 N-terminal domains of the *Cetobacterium somerae* prediction (UniProt U7V983) were kept and those with Cα RMSD < 4 Å to the resting-state cryo-EM structure or Cα RMSD < 3 Å to the reactivation state structure (PDB 3BUL) were excluded. Finally, the number of models in the reactivation state were determined by alignment to the 3BUL crystal structure (Cα RMSD < 2.5 Å).

**Supplementary Table 1: SEC-SAXS data collection parameters for *E. coli* MetH (2-1227)**

| **(a) Sample details** | | | |
|---|---|---|---|
| Sample | Cob(II) MetH | $CH_3$-Cob(III) MetH | As-isolated MetH |
| Source organism | *Escherichia coli* | *Escherichia coli* | *Escherichia coli* |
| Expression system | *Escherichia coli* | *Escherichia coli* | *Escherichia coli* |
| *M* from chemical composition (with cofactors) | 137263 | 137278 | ~137278 |
| Extinction coefficient (280 nm, theoretical) | 136,600 $M^{-1}$ $cm^{-1}$ | 136,600 $M^{-1}$ $cm^{-1}$ | 136,600 $M^{-1}$ $cm^{-1}$ |
| Loading concentration | 31 µM | 42 µM | 100 µM |
| Buffer composition | 50 mM HEPES, 150 mM NaCl, 2.5 mM DTT pH 7.6 | 50 mM HEPES, 150 mM NaCl, 2.5 mM DTT pH 7.6 | 50 mM HEPES, 150 mM NaCl, 2.5 mM DTT pH 7.6 |
| **(b) SAXS data collection parameters** | | | |
| Sample | Cob(II) MetH | $CH_3$-Cob(III) MetH | As-isolated MetH |
| Beamline/detector | CHESS ID7A/Pilatus 100K | CHESS ID7A/Pilatus 100K | CHESS ID7A/Pilatus 100K |
| Energy (keV) | 9.9 | 9.9 | 9.9 |
| Beam size (µm) | 250 × 250 | 250 × 250 | 250 × 250 |
| Detector distance (m) | 1.53 | 1.53 | 1.53 |
| *q*-measurement range ($Å^{-1}$) | 0.0091-0.275 | 0.0091-0.275 | 0.0091-0.275 |
| Normalization | Transmitted intensity | Transmitted intensity | Transmitted intensity |
| Exposures | 493 × 2 s | 587 × 2 s | 578 × 2 s |
| Configuration (batch/SEC) | SEC | SEC | SEC |
| Column (flow rate, ml/min) | Superdex 200 5/150 GL (0.2) | Superdex 200 5/150 GL (0.2) | Superdex 200 5/150 GL (0.2) |
| Experimental temperature (°C) | 4 | 4 | 4 |
| **(c) Data processing/structural parameters (main component)** | | | |
| Sample | Cob(II) MetH | $CH_3$-Cob(III) MetH | As-isolated MetH |
| Buffer Range (frames) | 92-146 | 95-154 | 49-100 |
| Main component range (frames) | 222-355 (EFA) | 230-380 (EFA) | 260-282 (average) |
| **Guinier Analysis** | | | |
| $I(0)$ | 3.78 ± 0.011 | 4.05 ± 0.0082 | 0.21 ± 0.00021 |
| $R_g$ (Å) | 41.27 ± 0.18 | 40.4 ± 0.12 | 40.19 ± 0.064 |
| $qR_g$ range | 0.374-1.263 | 0.366-1.282 | 0.385-1.297 |
| $R^2$ (fit) | 0.996 | 0.998 | 0.995 |
| ***P*(*r*) analysis (GNOM)** | | | |
| $I(0)$ | 3.81 ± 0.011 | 4.05 ± 0.0072 | 0.21 ± 0.0020 |
| $R_g$ (Å) | 41.99 ± 0.16 | 41.12 ± 0.08 | 40.38 ± 0.045 |
| $D_{max}$ (Å) | 147 | 134 | 128 |
| *q*-range ($Å^{-1}$) | 0.0091-0.275 | 0.0091-0.275 | 0.0091-0.275 |
| $\chi^2$ /total estimate | 1.19/0.74 | 1.34/0.96 | 1.20/0.91 |
| *MW*, kDa ($V_P$) | 146.3 | 150.5 | 134.6 |

**Supplementary Table 2: SAXS data collection parameters for *E. coli* MetH oxidation states**

| **(a) Sample details** | | | |
|---|---|---|---|
| Sample | Cob(I) MetH | Cob(II) MetH | $CH_3$-Cob(III) MetH |
| Source organism | *Escherichia coli* | *Escherichia coli* | *Escherichia coli* |
| Expression system | *Escherichia coli* | *Escherichia coli* | *Escherichia coli* |
| *M* from chemical composition (with cofactors) | 137263 | 137263 | 137278 |
| Extinction coefficient (280 nm, theoretical) | 136,600 $M^{-1}$ $cm^{-1}$ | 136,600 $M^{-1}$ $cm^{-1}$ | 136,600 $M^{-1}$ $cm^{-1}$/ 8910 $M^{-1}$ $cm^{-1}$ (525 nm, $B_{12}$) |
| Loading concentration | 20 µM | 20 µM | 20 µM |
| Buffer composition | 50 mM HEPES, 150 mM NaCl, 5 mM DTT pH 7.6 | 50 mM HEPES, 150 mM NaCl, 2.5 mM DTT pH 7.6 | 50 mM HEPES, 150 mM NaCl, 2.5 mM DTT pH 7.6 |
| **(b) SAXS data collection parameters** | | | |
| Sample | Cob(I) MetH | Cob(II) MetH | $CH_3$-Cob(III) MetH |
| Beamline/detector | CHESS ID7A/Eiger 4M | CHESS ID7A/Eiger 4M | CHESS ID7A/Eiger 4M |
| Energy (keV) | 10.1 | 14 | 10.1 |
| Beam size (µm) | 250 × 250 | 250 × 250 | 250 × 250 |
| Detector distance (m) | 1.60 | 1.71 | 1.60 |
| *q*-measurement range ($Å^{-1}$) | 0.0096-0.546 | 0.0082-0.529 | 0.0096-0.546 |
| Normalization | Transmitted intensity | Transmitted intensity | Transmitted intensity |
| Exposures | 20 × 1 s | 20 × 1 s | 20 × 1 s |
| Configuration (batch/SEC) | Batch (oscillating sample) | Batch (oscillating sample) | Batch (oscillating sample) |
| Experimental temperature (°C) | 20 | 20 | 20 |
| **(c) Data processing/structural parameters** | | | |
| Sample | Cob(I) MetH | Cob(II) MetH | $CH_3$-Cob(III) MetH |
| **Guinier Analysis** | | | |
| *I*(*0*) | 0.134 ± 0.00058 | 0.230 ± 0.00053 | 0.99 ± 0.0019 |
| $R_g$ (Å) | 42.06 ± 0.25 | 41.86 ± 0.14 | 40.01 ± 0.13 |
| $qR_g$ range | 0.404-1.293 | 0.342-1.292 | 0.385-1.297 |
| $R^2$ (fit) | 0.984 | 0.994 | 0.995 |
| ***P*(*r*) analysis (GNOM)** | | | |
| *I*(*0*) | 0.137 ± 0.00046 | 0.231 ± 0.00051 | 0.298 ± 0.00062 |
| $R_g$ (Å) | 43.43 ± 0.21 | 42.75 ± 0.11 | 40.99 ± 0.08 |
| $D_{max}$ (Å) | 157 | 152 | 140 |
| *q*-range ($Å^{-1}$) | 0.0096-0.546 | 0.0082-0.529 | 0.0096-0.546 |
| $\chi^2$ /total estimate | 1.06/0.805 | 1.12/0.781 | 1.02/0.873 |
| *MW*, kDa ($V_P$) | 142.1 | 138.9 | 138.4 |

**Supplementary Table 3: SAXS data collection parameters for $CH_3$-$H_4$folate titration series**

| **(a) Sample details** | | |
|---|---|---|
| Sample | Cob(II) MetH | |
| Source organism | *Escherichia coli* | |
| Expression system | *Escherichia coli* | |
| *M* from chemical composition (with cofactors) | 137263 | |
| Extinction coefficient (280 nm, theoretical) | 136,600 $M^{-1}$ $cm^{-1}$ | |
| Loading concentration | 20 µM | |
| Buffer composition | 50 mM HEPES, 150 mM NaCl, 2.5 mM DTT pH 7.6 | |
| Substrate (MW, g/mol) | (6*S*)-5-methyl-5,6,7,8-tetrahydropteroyl-n-γ-glutamic acid (445.4) | |
| Substrate concentrations (µM) | 0, 5, 10, 20, 50, 100, 500, 1000 | |
| **(b) Data collection parameters** | | |
| Beamline/detector | CHESS ID7A/Eiger 4M | |
| Energy (keV) | 14 | |
| Beam size (µm) | 250 × 250 | |
| Detector distance (m) | 1.71 | |
| *q*-measurement range ($Å^{-1}$) | 0.0082-0.529 | |
| Normalization | Transmitted intensity (beam-stop counter) | |
| Exposures | 20 × 1 s | |
| Configuration (batch/SEC) | Batch (oscillating sample) | |
| Experimental temperature (°C) | 20 | |
| **(c) Titration data statistics** | | |
| **$CH_3$-$H_4$folate concentration (mM)** | **$R_g$, Å (Guinier)** | **$I(0)$ (Guinier)** |
| 0 | 41.91 ± 0.15 | 0.230 |
| 5 | 42.18 ± 0.14 | 0.250 |
| 10 | 42.86 ± 0.15 | 0.226 |
| 20 | 42.88 ± 0.14 | 0.236 |
| *50* | 42.94 ± 0.15 | 0.235 |
| 100 | 42.95 ± 0.15 | 0.233 |
| 500 | 43.07 ± 0.15 | 0.243 |
| 1000 | 43.10 ± 0.14 | 0.238 |

**Supplementary Table 4: SAXS data collection parameters for Cob(II)/FldA titration series**

| **(a) Sample details** | | |
|---|---|---|
| Sample | Cob(II) MetH | FldA flavodoxin |
| Source organism | *Escherichia coli* | *Escherichia coli* |
| Expression system | *Escherichia coli* | *Escherichia coli* |
| *M* from chemical composition (with cofactors) | 137263 | 21016 |
| Extinction coefficient (280 nm, theoretical) | 136,600 $M^{-1}$ $cm^{-1}$ | 31,200 $M^{-1}$ $cm^{-1}$/8250 $M^{-1}$ $cm^{-1}$ (466 nm, ox. FMN) |
| Loading concentration | 20 µM | 5-60 µM |
| Buffer composition | 44 mM Na/KPi, 1 mM DTT pH 6.0, ionic strength = 0.15 | 44 mM Na/KPi, 1 mM DTT pH 6.0, ionic strength = 0.15 |

| **(b) Data collection parameters** | |
|---|---|
| Beamline/detector | CHESS ID7A/Eiger 4M |
| Energy (keV) | 14 |
| Beam size (µm) | 250 × 250 |
| Detector distance (m) | 1.71 |
| *q*-measurement range ($Å^{-1}$) | 0.0082-0.529 |
| Normalization | Transmitted intensity (beam-stop counter) |
| Exposures | 20 × 1 s |
| Configuration (batch/SEC) | Batch (oscillating sample) |
| Experimental temperature (°C) | 20 |

| **(c) Titration data statistics** | | |
|---|---|---|
| **FldA concentration (µM)** | **$R_g$, Å (Guinier)** | ***I*(*0*) (Guinier)** |
| 0 | 43.08 ± 0.18 | 0.242 |
| 5 | 44.35 ± 0.17 | 0.271 |
| 10 | 44.94 ± 0.26 | 0.290 |
| 20 | 45.06 ± 0.20 | 0.298 |
| 25 | 45.30 ± 0.18 | 0.322 |
| 30 | 45.18 ± 0.19 | 0.328 |
| 40 | 44.55 ± 0.23 | 0.341 |
| 60 | 43.22 ± 0.15 | 0.352 |
| 20 µM FldA only | 17.91 ± 0.11 | 0.207 |

**Supplementary Table 5: SAXS data collection parameters for Cob(III)/FldA titration series**

| **(a) Sample details** | | |
|---|---|---|
| Sample | $CH_3$-Cob(III) MetH | FldA flavodoxin |
| Source organism | *Escherichia coli* | *Escherichia coli* |
| Expression system | *Escherichia coli* | *Escherichia coli* |
| *M* from chemical composition (with cofactors) | 137278 | 21016 |
| Extinction coefficient (280 nm, theoretical) | 136,600 $M^{-1}$ $cm^{-1}$/8910 $M^{-1}$ $cm^{-1}$ (525 nm, $B_{12}$) | 31,200 $M^{-1}$ $cm^{-1}$/8250 $M^{-1}$ $cm^{-1}$ (oxidized FMN) |
| Loading concentration | 20 µM | 5-60 µM |
| Buffer composition | 44 mM Na/KPi, 1 mM DTT pH 6.0, ionic strength = 0.15 | 44 mM Na/KPi, 1 mM DTT pH 6.0, ionic strength = 0.15 |

| **(b) Data collection parameters** | |
|---|---|
| Beamline/detector | CHESS ID7A/Eiger 4M |
| Energy (keV) | 14 |
| Beam size (µm) | 250 × 250 |
| Detector distance (m) | 1.71 |
| *q*-measurement range (Å $^{-1}$) | 0.0082-0.529 |
| Normalization | Transmitted intensity (beam-stop counter) |
| Exposures | 20 × 1 s |
| Configuration (batch/SEC) | Batch (oscillating sample) |
| Experimental temperature (°C) | 20 |

| **(c) Titration data statistics** | | |
|---|---|---|
| **FldA concentration (µM)** | $R_g$, **Å (Guinier)** | ***I*(*0*) (Guinier)** |
| 0 | 41.17 ± 0.14 | 0.221 |
| 5 | 41.74 ± 0.14 | 0.231 |
| 10 | 41.35 ± 0.1 | 0.255 |
| 20 | 41.05 ± 0.17 | 0.307 |
| 25 | 40.91 ± 0.16 | 0.324 |
| 30 | 40.61 ± 0.20 | 0.334 |
| 60 | 36.97 ± 0.21 | 0.368 |
| 20 µM FldA only | 17.79 ± 0.14 | 0.181 |

**Supplementary Table 6: SAXS data collection parameters for *T. filiformis* MetH**

| **(a) Sample details** | | |
|---|---|---|
| Sample | Cob(II) MetH | Cob(II) MetH |
| Source organism | *Thermus filiformis* | *Thermus filiformis* |
| Expression system | *Escherichia coli* | *Escherichia coli* |
| *M* from chemical composition (with cofactors) | 132539 | 132539 |
| Extinction coefficient (280 nm, theoretical) | 123,580 $M^{-1}$ $cm^{-1}$ | 123,580 $M^{-1}$ $cm^{-1}$ |
| Loading concentration | 20 µM | 20 µM |
| Buffer composition | 50 mM HEPES, 150 mM NaCl 2.5 mM DTT pH 7.6 | 50 mM HEPES, 150 mM NaCl 2.5 mM DTT pH 7.6 |
| Substrate (MW, g/mol) | N/A | 1 mM (6*S*)-5-methyl-5,6,7,8-tet-rahydropteroyl-n-γ-glutamic acid (445.4) |

| **(b) Data collection parameters** | |
|---|---|
| Instrument/detector | Xenocs BioXolver (Pilatus 300K) |
| Energy (keV) | 8 |
| Detector distance (m) | 3.2 |
| *q*-measurement range ($Å^{-1}$) | 0.0146-0.439 |
| Normalization | Transmitted intensity (direct beam) |
| Exposures | 10 × 120 s |
| Configuration (batch/SEC) | Batch (stationary sample) |
| Experimental temperature (°C) | 20 |

| **(c) Data processing/structural parameters** | | |
|---|---|---|
| Sample | Cob(II) MetH | Cob(II) MetH + $CH_3$-$H_4$folate |
| **Guinier Analysis** | 0.034 ± 0.00046 | 0.026 ± 0.00064 |
| *I*(*0*) | 46.73 ± 0.83 | 48.39 ± 1.11 |
| $R_g$ (Å) | 0.731-1.285 | 0.757-1.278 |
| $qR_g$ range | 0.985 | 0.973 |
| $R^2$ (fit) | | |
| ***P*(*r*) analysis (GNOM)** | | |
| *I*(*0*) | 0.033 ± 0.00042 | 0.035 ± 0.00083 |
| $R_g$ (Å) | 47.29 ± 0.66 | 49.69 ± 0.64 |
| $D_{max}$ (Å) | 175 | 180 |
| *q*-range ($Å^{-1}$) | 0.0199-0.439 | 0.021-0.439 |
| $\chi^2$ /total estimate | 1.08/0.757 | 0.938/0.0.773 |
| *MW*, kDa ($V_P$) | 153.9 | 146.8 |

**Supplementary Table 7: Cryo-EM data collection and processing**

| **(a) Sample details** | |
|---|---|
| Sample | Cob(II) MetH |
| Source organism | *Thermus filiformis* |
| Expression system | *Escherichia coli* |
| Freezing method (cryogen) | FEI Vitrobot MK IV (ethane/propane) |
| *M* from chemical composition (with cofactors) | 132539 |
| Grid support | Quantifoil r1.2/1.3 copper mesh |
| Concentration | 2 µM |
| Buffer condition | 50 mM HEPES, 150 mM NaCl 2.5 mM DTT pH 7.6 |
| Additional additives | 2 µM horse spleen apoferritin |
| **(b) Data Collection and Processing Statistics** | |
| Microscope | FEI Titan Krios |
| Accelarating voltage (keV) | 300 keV |
| Defocus range (µm) | 0.8-2.5 µm |
| Energy Filter (eV) | 30 eV |
| Detector | Gatan K3 |
| Micrograph movies: collected (used) | 12,768 (9,656) |
| Frames per image stack | 50 |
| Physical Pixel size (Å) | 1.0691 |
| Total dose ($e^-$/Å$^2$) | 61.7 |
| Particle box size: pixels (Å) | 384 (410.5) |
| Particles: initial (final) | 9,994,870 (257,706) |
| B-factor (Guinier) | 112.4 |
| Consensus map resolution, FSC=0.143/0.5 (Å) | 3.60/4.13 |
| Local resolution range (Å) | 3.1 – 9.2 (75%) |

**Supplementary Table 8: Cryo-EM model**

| **Model and model/map statistics** | |
|---|---|
| Residues modeled (range) | 841 (37-877) |
| Atoms (no hydrogens) | 6541 |
| Ligands | 2 |
| Resolution, map/model FSC = 0.5 (Å) | 4.48 |
| MolProbity score | 2.64 |
| *Bonds (RMSD)* | |
| Length (Å) (# > 4$\sigma$) | 0.015 (30) |
| Angles (°) (# > 4$\sigma$) | 2.379 (71) |
| Clashscore | 13.55 |
| *Ramachandran plot (%)* | |
| Favored | 91.30 |
| Allowed | 8.70 |
| Outliers | 0.0 |
| Rotamer outliers (%) | 4.42 |
| Cβ outliers (%) | 1.17 |
| CaBLAM outliers (%) | 3.23 |
| Cis proline/general | 0.0/0.0 |
| Twisted proline/general | 0.0/0.0 |
| *ADP (B-factors)* | |
| Protein (min/max/mean) | 7.80/71.02/24.59 |
| Ligand (min/max/mean) | 2.00/25.69/9.44 |
| CC (mask) | 0.52 |
| CC (box) | 0.59 |
| CC (volume) | 0.54 |

**Supplementary Table 9: EOM fit statistics (*E. coli* MetH 2-1227)**

| **Resting state model** | ***Substrate-free Cob(II) MetH*** |
|---|---|
| Structures in starting pool | 10,000 |
| Fit $\chi^2$ value | 1.211 |
| $R_{flex}$ (random) / $R_{sigma}$ | 84.80% (~ 89.73%) / 1.50 |
| Final ensemble $R_g$ (Å) | 41.45 |
| Final ensemble $D_{max}$ (Å) | 131.0 |
| Representative structures in final ensemble | 2 |
| Corresponding plot | 5B |

**Supplementary Table 10: SAXS results for 3-domain *E. coli* MetH (2-896)**

| **(a) Sample details** | | |
|---|---|---|
| Sample | Cob(II) MetH (2-896) | Cob(II) MetH (2-896) |
| Source organism | *Escherichia coli* | *Escherichia coli* |
| Expression system | *Escherichia coli* | *Escherichia coli* |
| *M* from chemical composition (with cofactors) | 98181 | 98181 |
| Extinction coefficient (280 nm, theoretical) | 67,270 $M^{-1}$ $cm^{-1}$ | 67,270 $M^{-1}$ $cm^{-1}$ |
| Loading concentration | 23 µM | 23 µM |
| Buffer composition | 50 mM HEPES, 150 mM NaCl, 2.5 mM DTT pH 7.6 | 50 mM HEPES, 150 mM NaCl, 2.5 mM DTT pH 7.6 |
| Substrate (MW, g/mol) | N/A | 1.2 mM (6*R*,*S*)-5-methyl-5,6,7,8-tetrahydropteroyl-n-γ-glutamic acid (445.4) |

| **(b) Data collection parameters** | |
|---|---|
| Beamline/detector | CHESS ID7A/Eiger 4M |
| Energy (keV) | 9.9 |
| Beam size (µm) | 250 × 250 |
| Detector distance (m) | 1.71 |
| *q*-measurement range (Å $^{-1}$) | 0.0095-0.4806 |
| Normalization | Transmitted intensity (beam-stop counter) |
| Exposures | 12 × 1 s |
| Configuration (batch/SEC) | Batch (oscillating sample) |
| Experimental temperature (°C) | 4 |

| **(c) Data processing/structural parameters** | | |
|---|---|---|
| Sample | Cob(II) (2-896) | Cob(II) (2-896) + $CH_3$-$H_4$folate |
| **Guinier Analysis** | | |
| $I(0)$ | 0.152 ± 0.00027 | 0.157 ± 0.00034 |
| $R_g$ (Å) | 32.41 ± 0.08 | 35.63 ± 0.12 |
| $qR_g$ range | 0.355-1.290 | 0.408-1.242 |
| $R^2$ (fit) | 0.995 | 0.992 |
| ***P*(*r*) analysis (GNOM)** | | |
| $I(0)$ | 0.150 ± 0.00025 | 0.160 ± 0.00031 |
| $R_g$ (Å) | 32.7 ± 0.08 | 36.7 ± 0.12 |
| $D_{max}$ (Å) | 122 | 143 |
| *q*-range (Å$^{-1}$) | 0.011-0.4806 | 0.0115-0.4806 |
| $\chi^2$ /total estimate | 1.063/0.828 | 1.159/0.731 |
| *MW*, kDa ($V_P$) | 109.2 | 107.0 |

**Supplementary Table 11: CRYSOL fit statistics (*E. coli* MetH 2-896)**

| **(a) Resting state model** | ***Substrate-free (2-896)*** | ***+ 0.6 mM $CH_3$-$H_4$folate (2-896)*** |
|---|---|---|
| Constant subtraction allowed | Yes | Yes |
| Spherical harmonics | 99 | 99 |
| Fit $\chi^2$ value | 1.488 | 7.721 |
| Experimental/Predicted $R_g$ (Å) | 32.41 / 31.93 | 35.63 / 31.29 |
| Corresponding plot | 6E/S18A | S18D |
| **(b) Homocysteine state model** | | |
| Constant subtraction allowed | Yes | Yes |
| Spherical harmonics | 99 | 99 |
| Fit $\chi^2$ value | 1.297 | 5.992 |
| Experimental/Predicted $R_g$ (Å) | 32.41 / 31.98 | 35.63 / 31.34 |
| Corresponding plot | S18B | S18E |
| **(c) Folate state model** | | |
| Constant subtraction allowed | Yes | Yes |
| Spherical harmonics | 99 | 99 |
| Fit $\chi^2$ value | 5.611 | 1.542 |
| Experimental/Predicted $R_g$ (Å) | 32.41 / 35.09 | 35.63 / 34.52 |
| Corresponding plot | S18C | 6E/S18F |

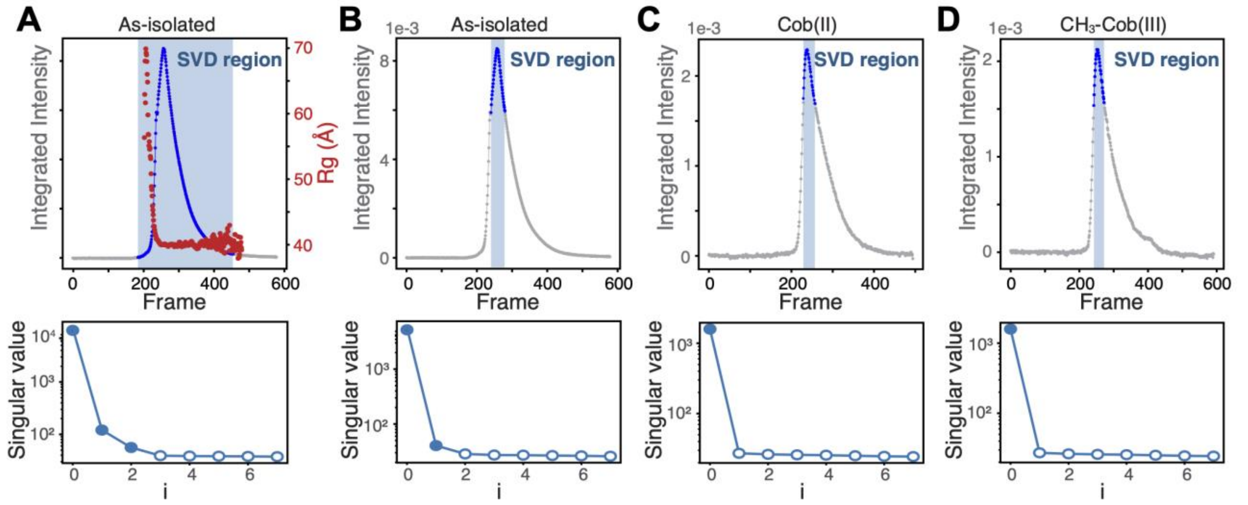


**Figure S1: SEC-SAXS of full-length *E. coli* MetH indicates the as-isolated protein is a mixture of multiple dominant conformations. (A)** SEC-SAXS of as-isolated full-length MetH produces a single elution peak (top). However, SVD reveals that the elution peak contains at least 3 major components (bottom). **(B)** Repeating SVD on a limited region (~50 frames near the top of the peak) shows that the main peak contains at least two, highly overlapping species that could not be computationally separated by deconvolution methods. By contrast, when similar SVD analysis is performed on the SEC-SAXS datasets of MetH prepared in the **(C)** Cob(II) state or **(D)** $CH_3$-Cob(III) state, the heterogeneity across the main peak disappears. This result suggests that when MetH is prepared in a single oxidation state, it is described by a single predominant conformation. Data processing statistics can be found in Table S1.

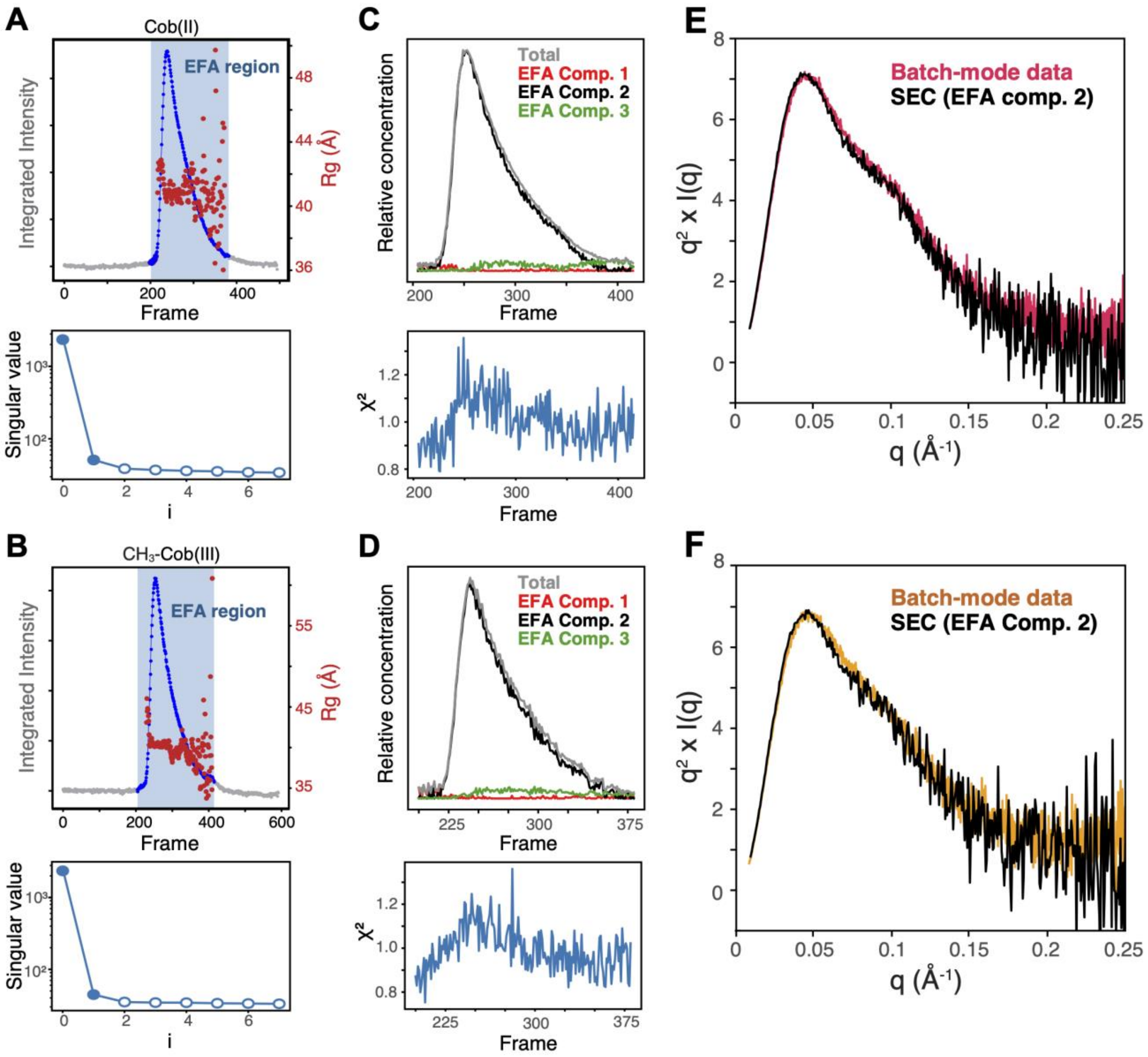


**Figure S2: SEC-SAXS of purified *E. coli* MetH oxidation states. (A)** SEC-SAXS of full-length Cob(II) MetH shown in Figure S1C. SVD of the entire elution peak (blue shaded region) indicates the presence of a minor second component. **(B)** SEC-SAXS of full-length $CH_3$-Cob(III) MetH shown in Figure S1D. SVD of the entire elution peak (blue shaded region) indicates the presence of a minor second component. **(C-D)** Deconvolution by evolving factor analysis (EFA)[11] of the blue shaded region shows that both Cob(II) and $CH_3$-Cob(III) MetH can be described predominantly as a single species (black) with very minor populations of a larger component (red) and a smaller component (green). The relative concentration peaks are normalized by the *I*(0) values of each component; the predominant component constitutes virtually all of the contribution to the scattering. **(E-F)** Comparison of the scattering from the dominant species in panels C and D are superimposable with our batch-mode data on Cob(II) and $CH_3$-Cob(III) MetH (first shown in Figure 2B), validating the use of batch-mode experiments for further analysis. Data processing statistics can be found in Table S1.

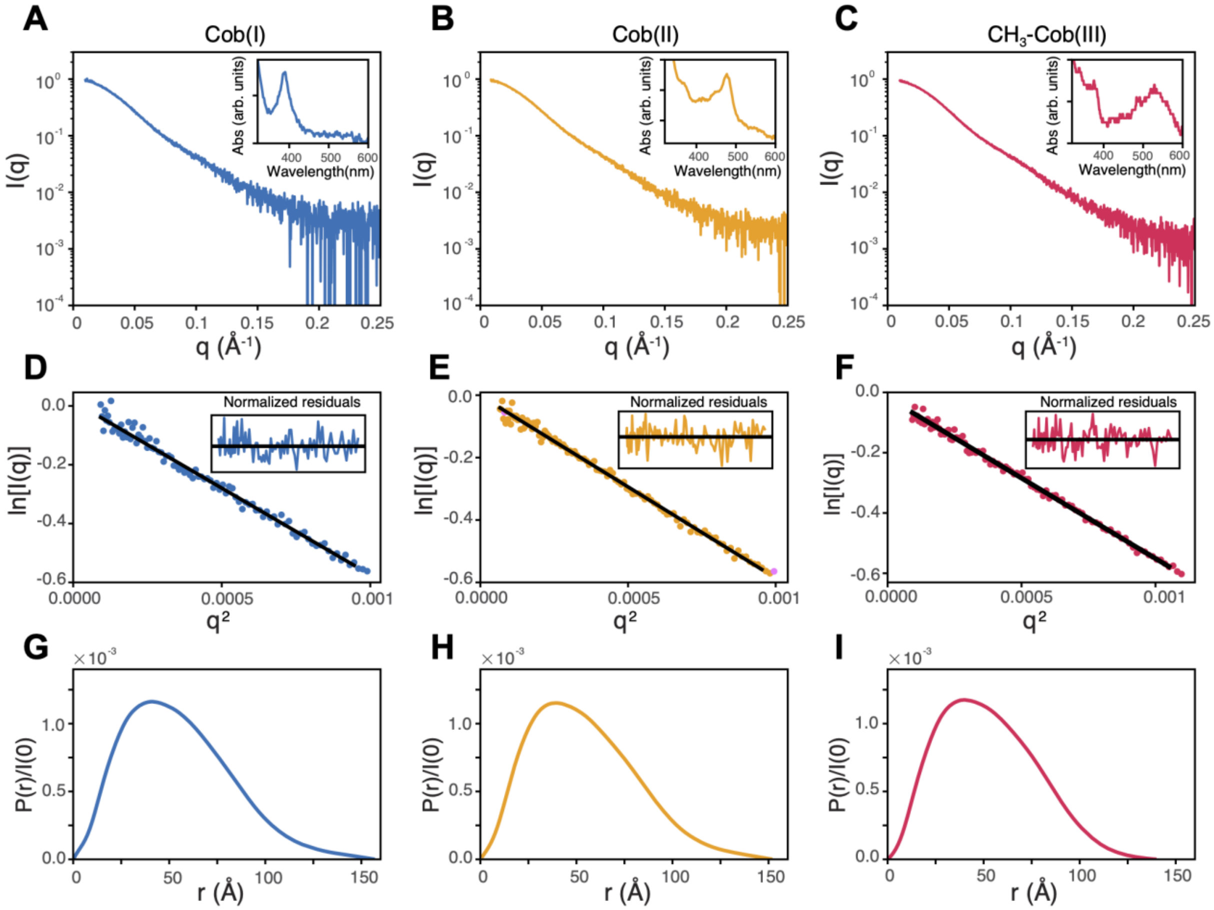


**Figure S3: Validation of batch-mode SAXS data for purified *E. coli* MetH oxidation states. (A-C)** SAXS data presented as semilog plots for pure Cob(I) (blue, panel A**)**, Cob(II) (orange, panel B) and $CH_3$-Cob(III) (red, panel C) MetH are the same data shown in Figure 2B. Insets: Respective UV-Vis absorption spectra taken post-exposure at the beamline agree well with the post-preparation spectra in Figure 2A. **(D-F)** Guinier fits for the SAXS data presented in panels A-C. Insets: The normalized residuals for the respective fits. All three datasets have good Guinier regions, without deviation from linearity at low-*q*. **(G-I)**. *P*(*r*) functions (calculated in GNOM[34]) for the SAXS data presented in panels A-C. The shapes of the *P*(*r*) functions for all three datasets are highly similar; the slightly longer tails (and resulting higher $D_{max}$ values) for the Cob(I) and Cob(II) datasets are likely a result of slightly degraded sample quality from the chemical preparation of those states from $CH_3$-Cob(III) enzyme. Data processing statistics can be found in Table S2.

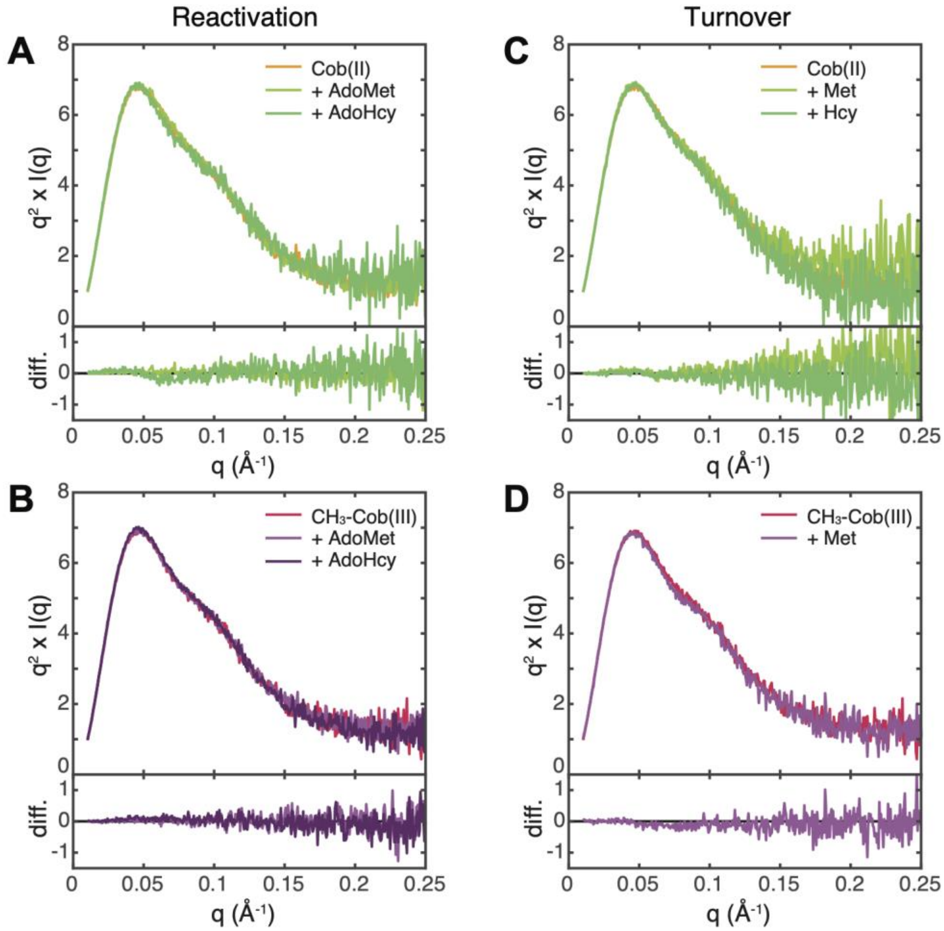


**Figure S4: Additional batch-mode SAXS experiments on full-length *E. coli* MetH with substrates and products from the reactivation and turnover cycles.** Incubation of 20 μM *E. coli* MetH with 1 mM *S*-adenosylmethionine (AdoMet) or *S*-adenosylhomocysteine (AdoHcy), appears to have no discernable effect on the conformation of either **(A)** Cob(II) or **(B)** $CH_3$-Cob(III) MetH. Likewise, incubation of 20 μM *E. coli* MetH with 1 mM L-homocysteine (Hcy) or L-methionine (Met) appears to have no discernable effect on the conformation ensemble of either **(C)** Cob(II) or **(D)** $CH_3$-Cob(III). The latter was incubated with Met only, as Hcy would turn over. SAXS data are presented as Kratky representations.

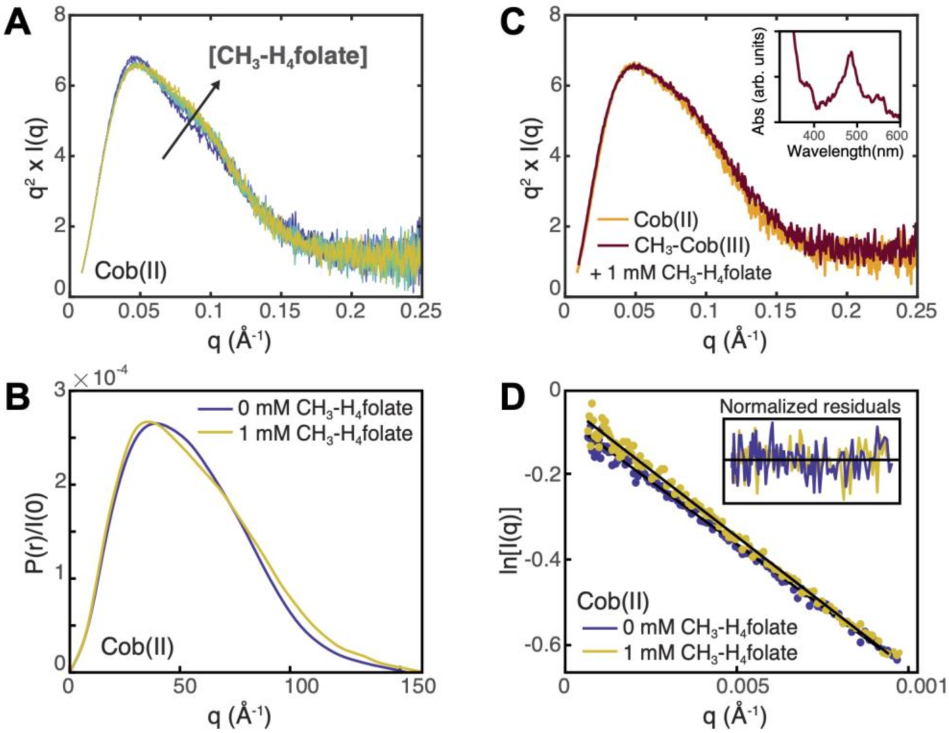


**Figure S5: Validation of batch-mode SAXS data for *E. coli* MetH folate titration series. (A)** Addition of 0-1 mM $CH_3$-$H_4$folate to 20 μM *E. coli* MetH in the Cob(II) state leads to a widening of the Kratky peak, indicative of a change to a slightly more elongated conformation. **(B)**. *P*(*r*) curves (calculated in GNOM[34]) for the two endpoints of the titration series (shown in Figure 2D), i.e. without substrate (purple) and after incubation with 1 mM $CH_3$-$H_4$folate (orange). Although the maximum dimension ($D_{max}$) does not change, the change in the shape of the *P*(*r*) curve suggests that MetH adopts a more elongated, rod-like shape with $CH_3$-$H_4$folate. **(C)** Addition of 1 mM $CH_3$-$H_4$folate to 20 μM *E. coli* MetH in $CH_3$-Cob(III) state (dark red) produces an identical conformational change as that observed for Cob(II) enzyme (orange). Interestingly, X-ray exposure unavoidably converted the enzyme to the Cob(II) state with an absorption maximum at 477 nm (inset), suggesting that the $CH_3$-$H_4$folate-driven conformational change involves the uncapping of the $B_{12}$ domain. **(D)** Guinier fits for the titration endpoints for Cob(II) *E. coli* MetH (shown in panel B). Insets are the normalized residuals for the respective fits. Both datasets have good Guinier regions, without deviation from linearity at low-*q*. Data processing statistics can be found in Table S3.

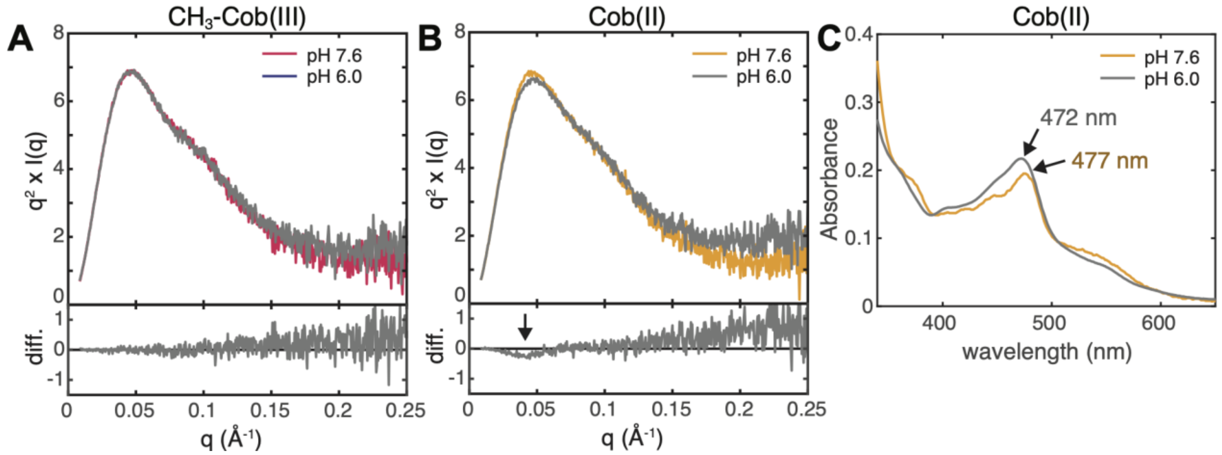


**Figure S6: pH dependence of *E. coli* MetH. (A)** The Kratky curves for full-length $CH_3$-Cob(III) MetH at pH 7.6 and 6.0 are superimposable. At both pH values, $CH_3$-Cob(III) MetH is predominantly His-on[35,36]. **(B)** The Kratky curves for full-length Cob(II) MetH at pH 7.6 and 6.0 are highly similar, although a subtle difference can be seen at $q$~0.045 Å$^{-1}$ (arrow). **(C)** The slight differences in scattering seen in panel B are potentially due to a higher population of His-off Cob(II) enzyme at lower pH, which is supported by a shift in the characteristic UV-Vis absorbance peak from ~477 nm (at pH 7.6) to ~472 nm (at pH 6.0). His-off Cob(II) MetH has an absorption maximum at ~465 nm[37].

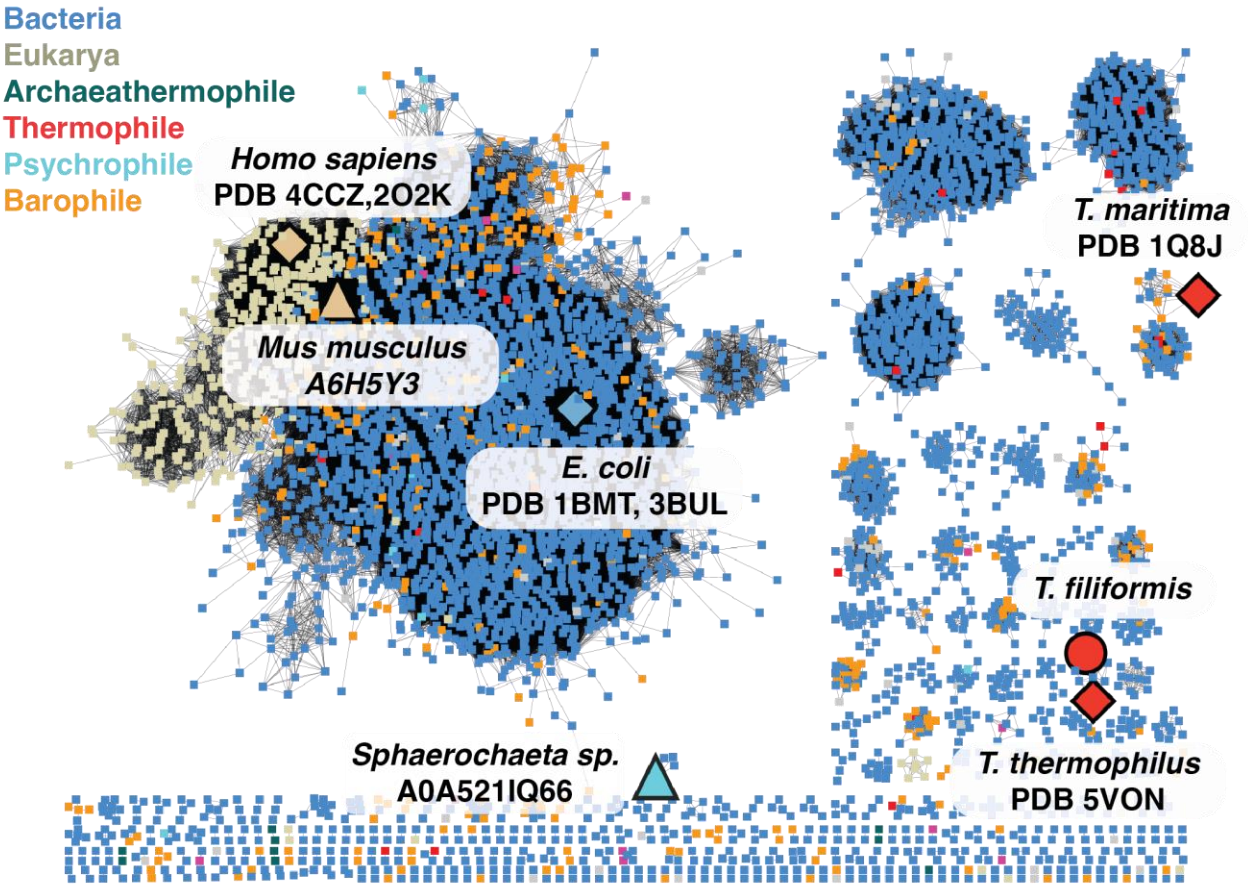


**Figure S7: MetH sequence-similarity network.** Visualization of the sequence-similarity network for cobalamin-dependent methionine synthase (Pfam PF02965), colored by organism classification: eukarya (tan), bacteria (blue), archaeathermophile (dark green), thermophile (red), psychrophile (light blue), barophile (>50 m depth, orange). Species with prior PDB depositions are marked by diamonds, species where the AlphaFold2 models were shown in Figure 6 are marked by triangles, and the species of which the cryo-EM structure was solved in this work is marked by a circle.

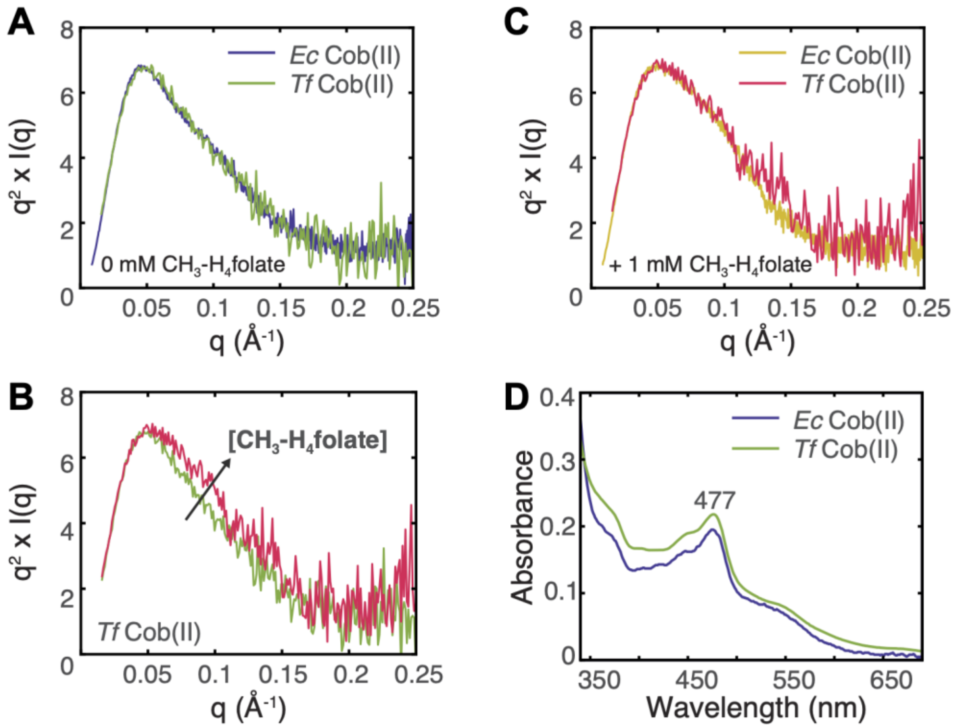


**Figure S8: SAXS suggests that *T. filiformis* MetH behaves similarly in solution to the *E. coli* enzyme. (A)** Comparison of 20 μM *T. filiformis* MetH (green) and *E. coli* MetH (blue) in the Cob(II) state in the absence of substrates. The Kratky curves are superimposable. **(B)** The Kratky curves for 20 μM *T. filiformis* MetH (red) and *E. coli* MetH (orange) in the presence of 1 mM $CH_3$-$H_4$folate are also superimposable. **(C)** The two Kratky curves for the *T. filiformis* enzyme shown in panels A-B. A qualitatively similar conformational change is observed upon addition of $CH_3$-$H_4$folate to the *T. filiformis* enzyme as was observed with the *E. coli* enzyme in Figure 2E. **(D)** UV-Vis spectra for substrate-free Cob(II) *E. coli* and *T. filiformis* MetH agree overall in shape and peak position, indicating that both are primarily His-on.

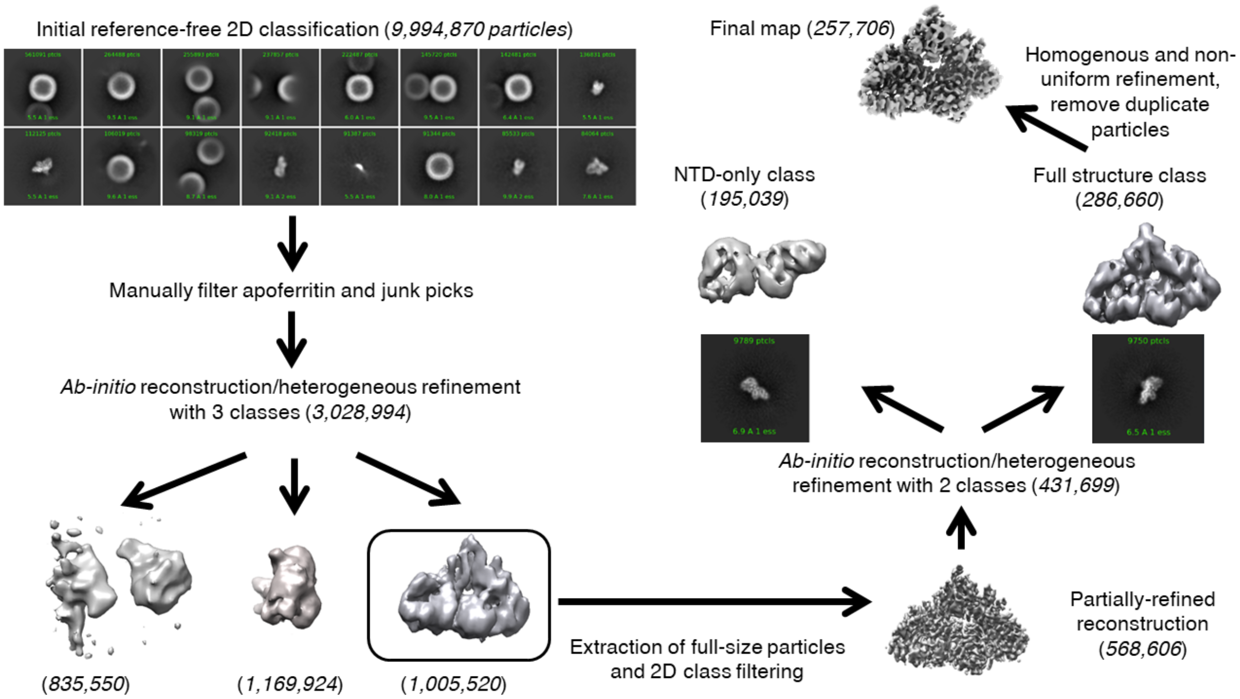


**Figure S9: Visualization of the cryo-EM data processing pipeline leading to the *T. filiformis* MetH consensus reconstruction**. An initial set of ~10 million particles was picked from 9,656 micrographs. Most of these particles represented apoferritin picks and were readily filtered out following initial reference-free 2D classification. The remaining particles were classified into three classes, representing a junk or highly heterogeneous class, a class containing just the two N-terminal domains, and a class containing the structure of interest. Following extraction of the full-size particles and further filtering by reference-free 2D classification, the remaining particle stack was separated into two classes, with one reconstruction containing just the two N-terminal domains and the other containing the N-terminal domains and the $B_{12}$ domain. The latter class was further trimmed by 2D classification and removal of duplicate particles and followed by homogenous and non-uniform refinement (without CTF or defocus refinement) to yield the final 3.6 Å map from 257,706 particles. The entire processing pipeline was performed in cryoSPARC[27]. Data processing statistics can be found in Table S7.

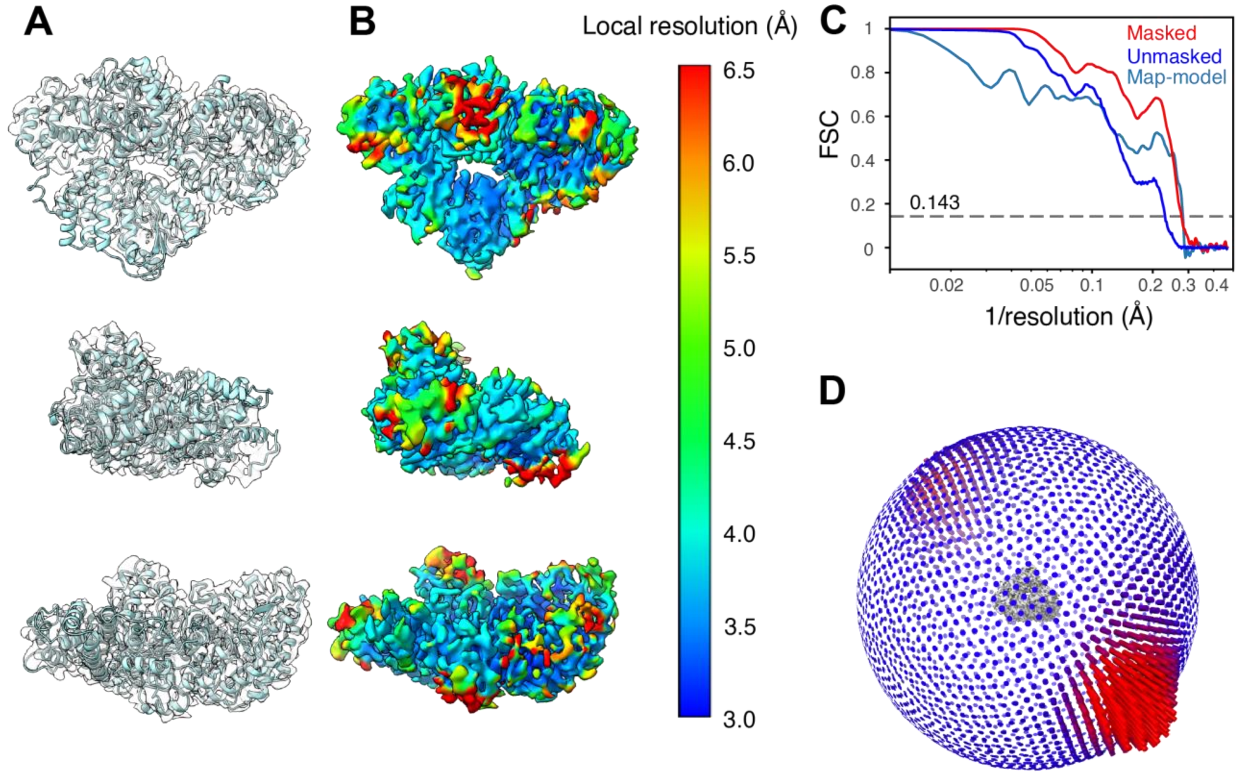


**Figure S10: Map and model validation for the *T. filiformis* cryo-EM consensus reconstruction. (A)** Orthogonal views of the model built into the consensus map. **(B)** Orthogonal map views, plotted by local resolution, in Å. The resolution of the majority of the map varies from ~3.3-6.5 Å and is generally much better at the cores of the domains. Local resolution maps were calculated in cryoSPARC[27]. **(C)** Masked and unmasked half-map Fourier Shell Correlation (FSC) curves for the consensus reconstruction, as well as the model-map FSC curve. The overall resolution of the map is 3.60 Å by FSC=0.143, although the interpretable level of detail is for most of the map closer to the 4-4.5 Å range. The map exhibits some resolution anisotropy likely due to remaining orientation bias. FSC curves were calculated in EMAN2[38]. **(D)** Distribution of observed views in the final particle stack. There are two dominant views; the orientation bias is strong enough to introduce some resolution anisotropy but not enough to cause problematic artifacts (i.e., incorrect dimensionality along an axis) in the final map as the dataset was large enough to sample other views. EM maps are shown at a threshold level of 0.3. Map and model statistics can be found in Tables S7 and S8.

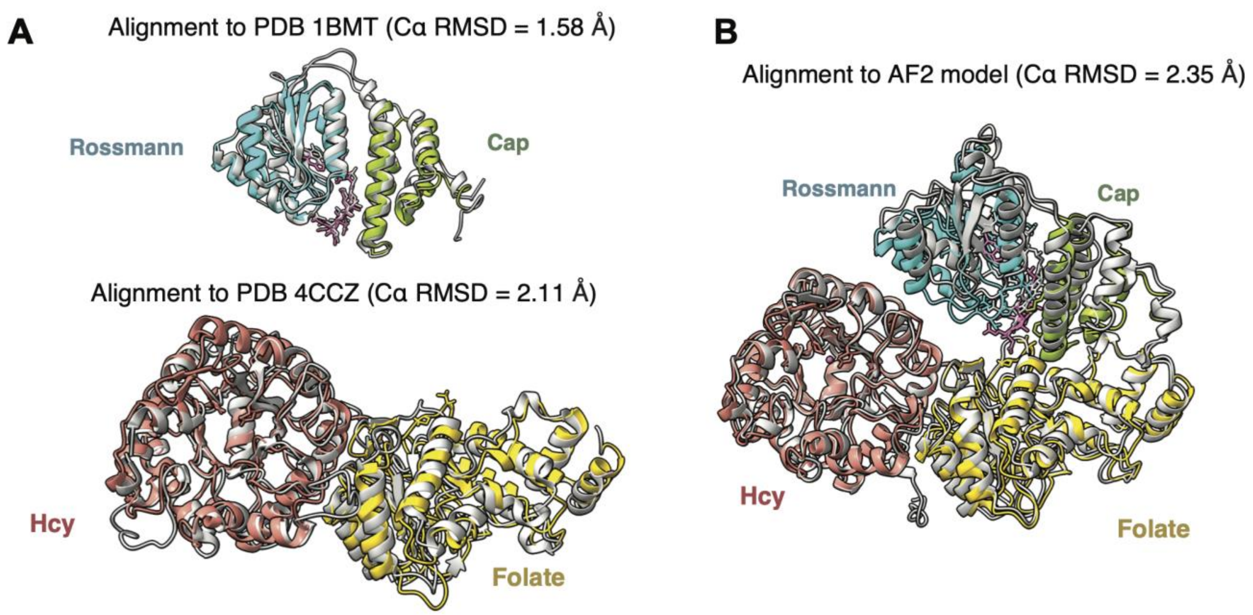


**Figure S11: Comparison of the *T. filiformis* cryo-EM model conformations to observed and predicted MetH structures. (A)** Sub-conformations of our cryo-EM consensus model are similar to previous crystal structures of MetH fragments. Top: The conformation adopted by the $B_{12}$-subdomains in our structure (colored) is highly similar to the "cap-on" conformation observed in the crystal structure of the *E. coli* $B_{12}$ domain alone (1BMT)[33] (light gray). Bottom: The overall orientation of the N-terminal domains in our structure (colored) is highly similar to that observed in the unpublished crystal structure of the two N-terminal domains from human MetH (4CCZ) (light gray). **(B)** The publicly available AlphaFold2[19] model for *T. filiformis* MetH (light gray) is remarkably similar to the novel domain arrangement observed in our cryo-EM model (colored), despite not containing the cobalamin ligand. The AlphaFold2 model is shown without the AdoMet domain for clarity as it does not interact with any other part of the model. The cryo-EM model is colored by domain assignment as shown in Figure 4.

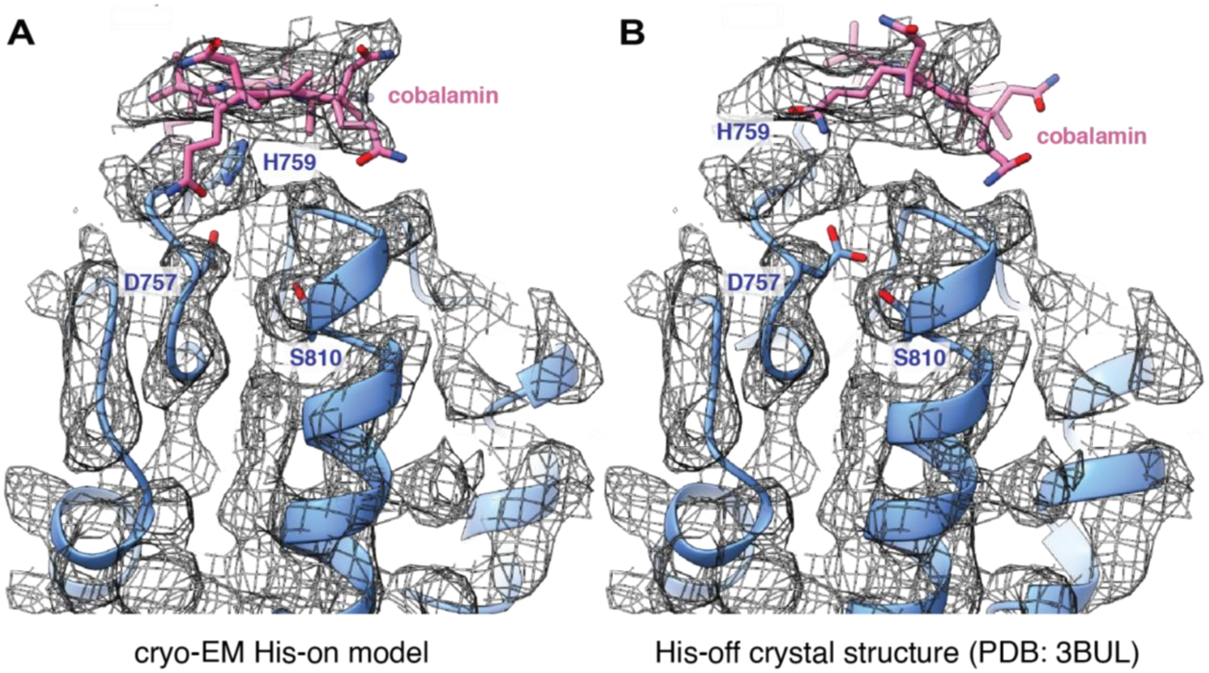


**Figure S12: The cryo-EM model supports a His-on cobalamin state. (A)** In our EM model of Cob(II) *T. filiformis* MetH, the orientation of the cobalamin that best fits the density is consistent with its position in the 1BMT[33] structure of the $B_{12}$ domain fragment, which is His-on. In our cryo-EM map (mesh) weak connecting density is visible between the central Co atom of cobalamin and His759, and Asp757 also appears to be in hydrogen bonding distance of His759, consistent with a His-on state. **(B)** When the His-off $B_{12}$ domain from the reactivation-state structure of *E. coli* MetH (3BUL[39]) is fit into the EM map, it is clear that the plane of the corrin ring as well as the orientation of Asp757 are shifted to positions not consistent with our map. EM maps are shown at a threshold level of 0.3. Note that by coincidence, Asp757, His759 and Ser810 have the same residue numbering in the *T. filiformis* and *E. coli* sequences.

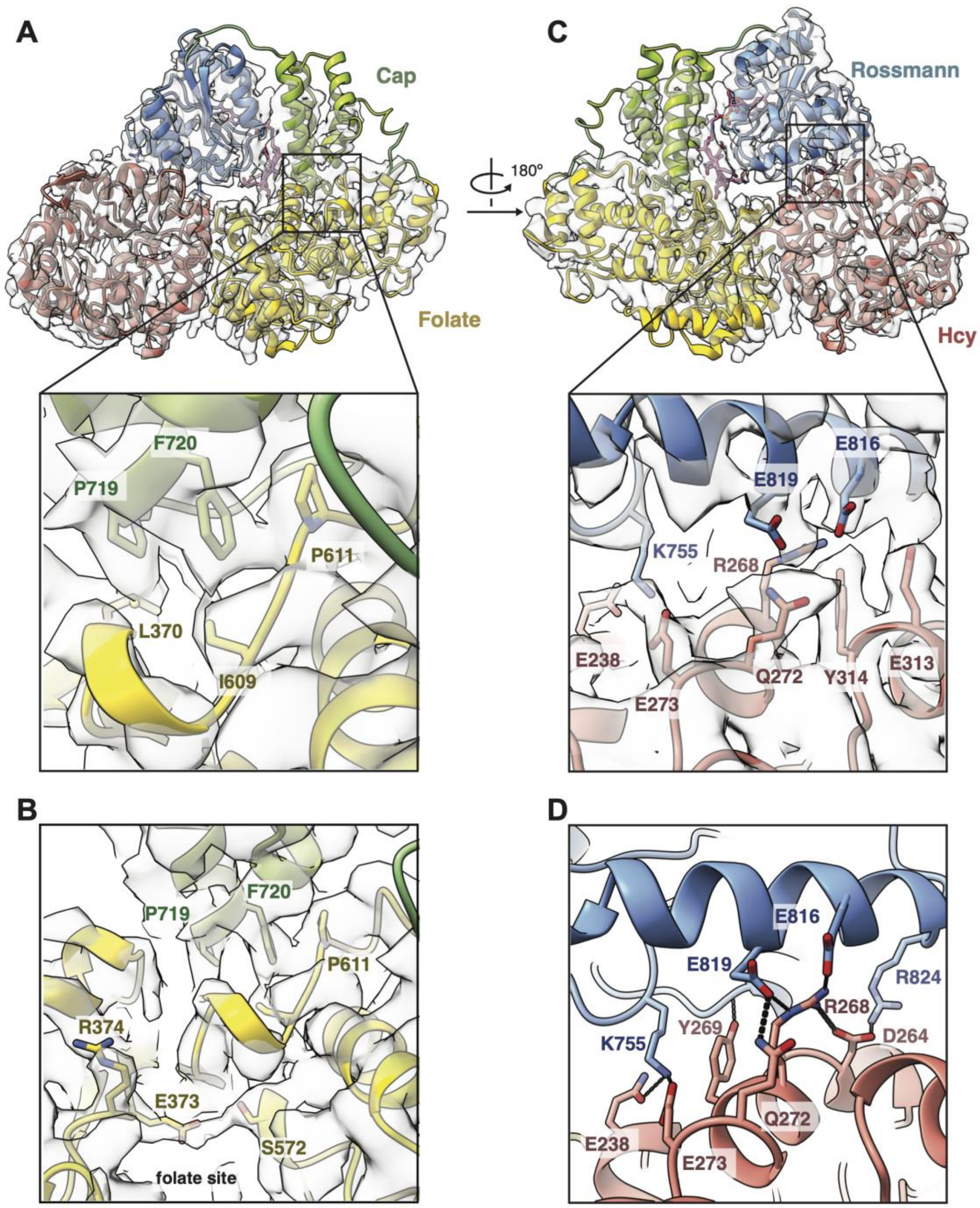


**Figure S13: Interactions between the $B_{12}$ domain and the N-terminal domains in the *T. filiformis* MetH resting-state structure. (A)** A hydrophobic patch interaction is observed between the cap subdomain (green) and the folate domain (yellow), including an aromatic-proline stacking interaction between P611 (on the folate domain) and F720 (on the cap subdomain). **(B)** This interaction appears to push the loop containing P611 towards the folate-binding site than in previous structures[40] as shown in Figure 4D-E. This loop adopts a different conformation when $CH_3$-$H_4$folate is bound. **(C-D)** The interaction between the Rossmann subdomain (blue) and the Hcy domain (salmon) consists of an extensive network of charged and polar residues. In particular, Asp264, Arg268, Gln272, Tyr269, and Glu273 of the α6-helix in the Hcy domain makes a number of polar interactions with the Rossmann subdomain. Density is somewhat inconsistent especially on the $B_{12}$ subdomain but does support the modeled positions of most of the putative interacting residues on the Hcy domain.

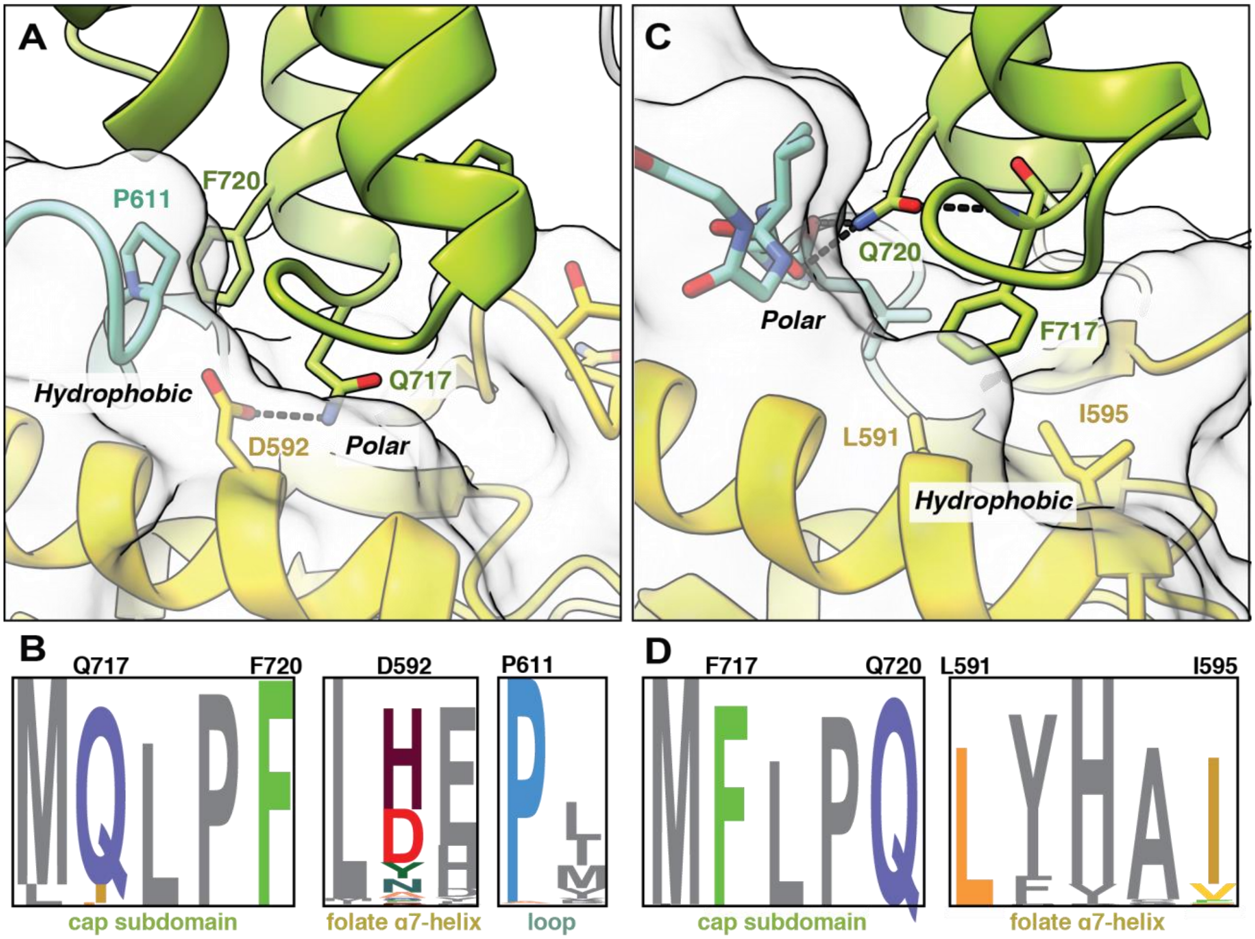


**Figure S14: AlphaFold2 analysis suggests that interdomain interactions are conserved in MetH from different species. (A)** In our cryo-EM structure, Phe720 on the cap subdomain (green) is observed making an aromatic-proline stacking interaction with Pro611 in the folate-binding loop (cyan) of the folate domain (yellow), while Gln717 appears to participate in a hydrogen-bonding interaction with Asp592 in the α7-helix of the folate domain (yellow). **(B)** Sequence logos of all MetH sequences in our SSN with a Phe at position 720 (*T. filiformis* numbering) show that the majority contain a Gln at position 717, a Pro at position 611, and a residue that can form polar interactions with Gln at position 592. **(C)** In E. coli MetH, the Phe and Gln positions flipped. Although there are no resting-state AlphaFold2[19] predictions of *E. coli* MetH, there is one for *Serinicoccus sp*. (Uniprot A0A1Q9SZ17, shown here in *T. filiformis* numbering), which is highly similar to the *E. coli* enzyme in this region. Gln720 in the cap subdomain (green) is observed making hydrogen-bonding interactions with the backbone carbonyl oxygens in the folate-sensing loop (cyan) as well as with the backbone amide of Phe717. Additionally, in place of the hydrogen-bonding interaction observed in our cryo-EM structure, here, we observe Leu591/Ile595 on the folate domain (yellow) creating a hydrophobic pocket for Phe717 on the cap subdomain (green). **(D)** Sequence logos of all MetH sequences in our SSN with a Gln at position 720 (*T. filiformis* numbering) show that the majority contain a Phe at position 717, Leu at position 591, and Ile at position 595.

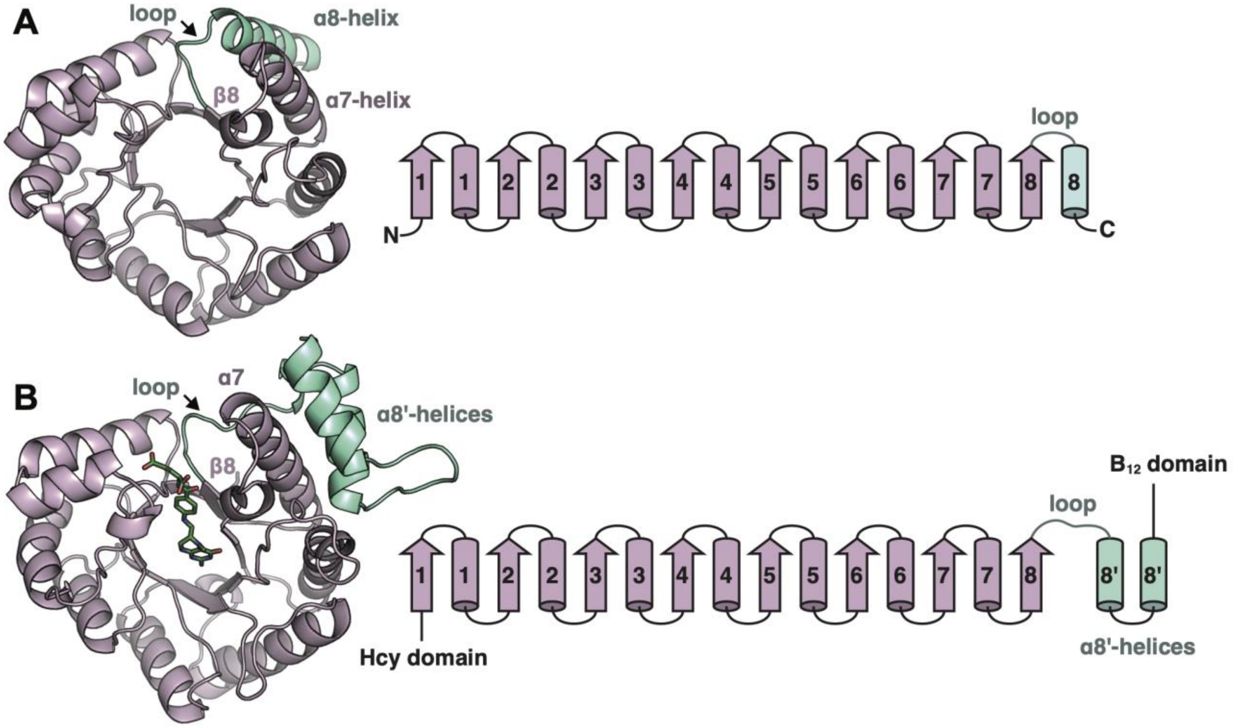


**Figure S15: Structure and topology of the MetH folate domain. (A)** A typical TIM barrel motif consists of a $\beta_8\alpha_8$ topology, where the loop extending from the β8-strand connects with the α8-helix, which completes the barrel structure. The structure of the N-terminal fragment of *Thermotoga maritima* MetH (PDB: 1Q8J) displays a canonical TIM barrel. However, this was attributed to the construct being truncated early[40]. **(B)** In all other MetH structures, including our own, the folate domain adopts a $\beta_8\alpha_7$ topology, where the loop from the β8-strand extends out to connect to a set of α8′-helices outside of the barrel. This was first observed in the structure of *Thermus thermophilus* MetH folate domain[40] (shown here).

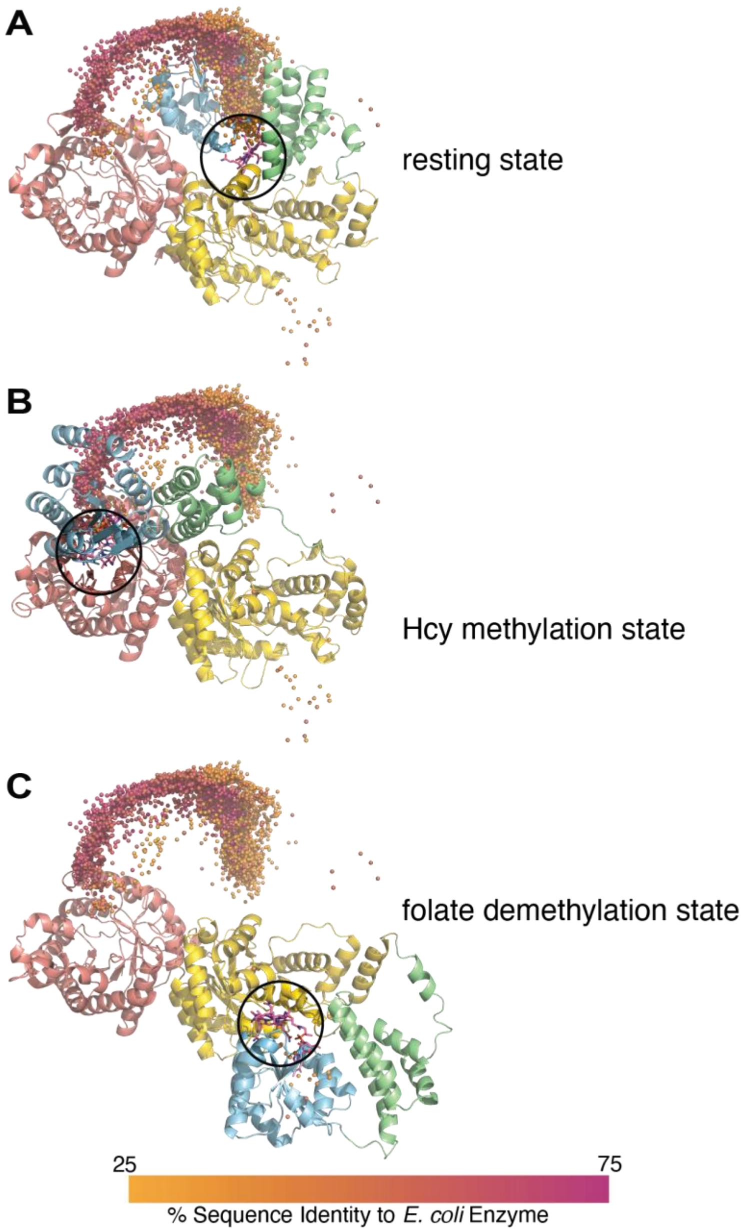


**Figure S16: Analysis of all 4-domain MetH sequences in the AlphaFold2 database.** Predicted models of 4915 MetH sequences (having a sequence length of 1100-1300 residues) contained in both the SSN (Figure S7) and the AlphaFold2 database were analyzed. The average positions of $B_{12}$-binding residues (as annotated in Uniprot) are visualized as dots. The terminal positions of these dots correspond to conformations that may have functional relevance (circled in black). Representative structures for each are shown in cartoon (without the AdoMet domain). **(A)** 212 models are predicted to be in the resting-state conformation observed by cryo-EM (shown). **(B)** 187 models are predicted to be in a homocysteine-methylation conformation (shown is the *Proteus mirabilis* model, Uniprot ID: A0A379GCZ0). **(C)** 6 models are predicted to be in a folate-demethylation conformation (shown is the *Cetobacterium somerae* model, Uniprot ID: U7V983). A continuum of dots is observed connecting the $B_{12}$ domain positions in the resting state and homocysteine-methylation state; interestingly, the $B_{12}$ domains in these models do not align well with the cap-on structure (1BMT[33]), suggesting that they are at least partially cap-off. AdoMet domain positions (not shown) for models that are not in the reactivation conformation generally have low confidence. The cobalamin ligand has been docked into the models for clarity.

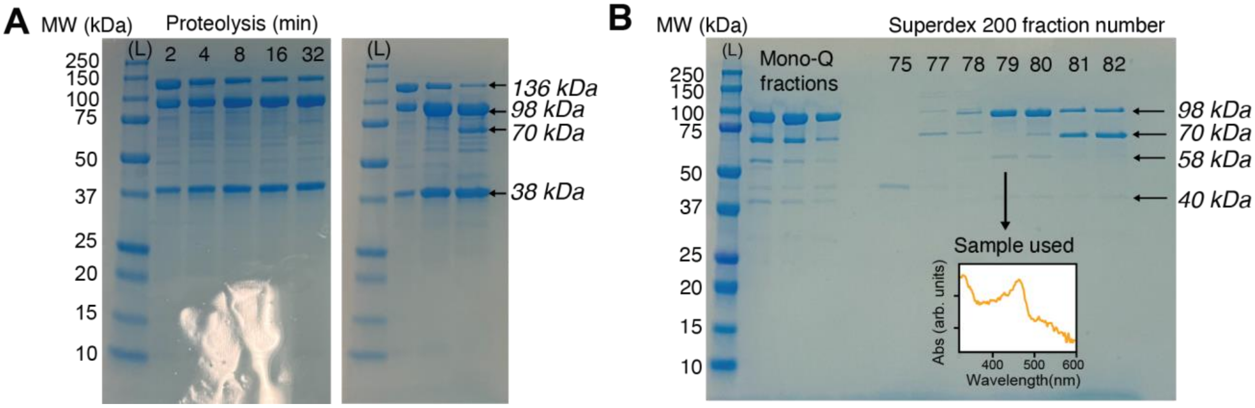


**Figure S17: Preparation of 3-domain *E. coli* MetH by limited proteolysis. (A)** Left: SDS-PAGE gel showing a time course of trypsin proteolysis of 30 µM full-length Cob(II) MetH, incubated with trypsin (final concentration 0.005 mg/mL) for 2-32 min. The leftmost lane is the ladder (L). Right: For the purification of the 3-domain fragment, 24 µM full-length Cob(II) MetH was incubated with trypsin at a final concentration 0.005 mg/mL. In the two lanes to the right of the ladder (L), the samples from 2- and 32-min time points from the time course test were run again as a control. The rightmost lane is the final sample used for purification (32 min incubation). The expected[10] 38 kDa (AdoMet domain) and 98 kDa (3 N-terminal domains) fragments are observed. The 38-kDa fragment and remaining undigested 136 kDa full-length protein were readily separated from the remaining species by anion exchange on a Mono-Q 5/150 column, but additional bands at 70, 58 and 40 kDa were also weakly present in the sample prior to loading onto the Mono-Q column, indicating some further proteolysis. **(B)** The left lanes show the fractions from the Mono-Q step that were loaded onto the SEC column. A very small amount of 58-kDa fragment is still present in these fractions. Subsequent size-exclusion purification separated the remaining unwanted fragments from the 98 kDa 3-domain fragment. Sample used for SAXS experiments came exclusively from a single SEC fraction, fraction 79. Inset: UV-Vis absorption spectroscopy of the purified 3-domain fragment confirmed the integrity of oxidation state of the cofactor in the His-on Cob(II) state. Overall yield of the preparation was ~13%.

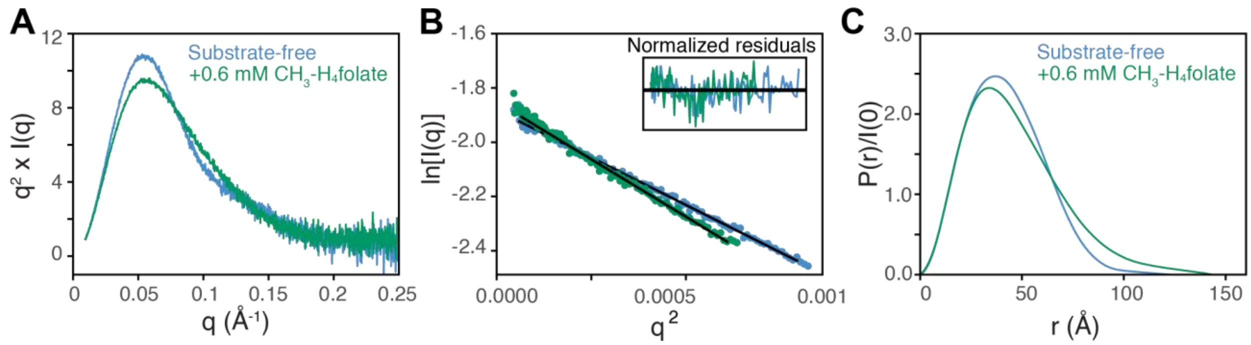


**Figure S18: Validation of batch-mode SAXS data for the 3-domain fragment of *E. coli* Cob(II) MetH (A)** SAXS data, presented in Kratky representation, for 24 μM 3-domain Cob(II) MetH without substrate (blue) and after incubation with 0.6 mM $CH_3$-$H_4$folate (green). Addition of $CH_3$-$H_4$folate leads to a widening of the Kratky peak, indicative of a change to a more elongated apparent state. (**B)** Guinier fits for the datasets shown in panel A with the same coloring. Inset is the normalized residuals for the respective fits. Both datasets have acceptable Guinier regions, without significant deviation from linearity at low-*q*. **(C)** *P*(*r*) curves (calculated in GNOM[34]) corresponding to the data shown in A. Without the AdoMet domain, it is readily apparent that the $CH_3$-$H_4$folate-incubated dataset represents a more elongated conformation. Data processing statistics can be found in Table S10.

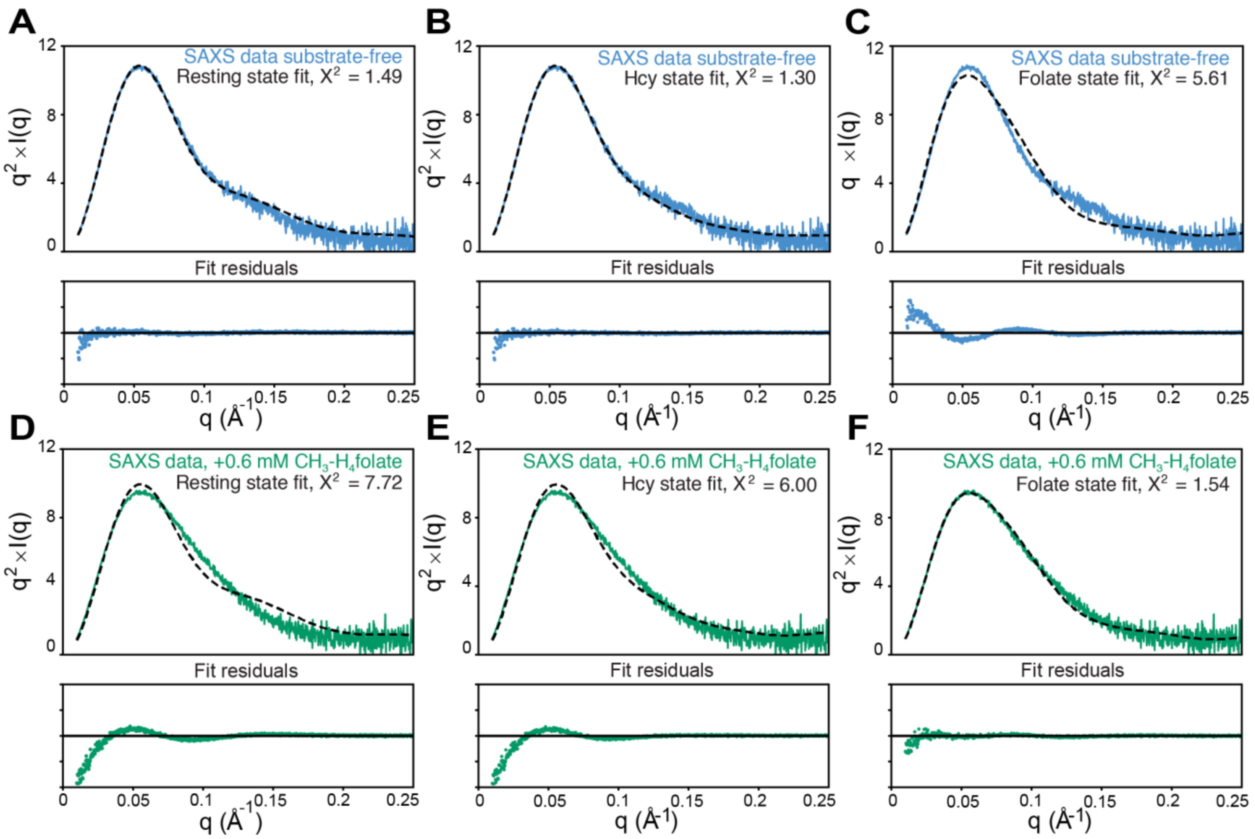


**Figure S19: Model fits to SAXS data on the 3-domain fragment of *E. coli* MetH.** Fits to SAXS data collected on 3-domain Cob(II) MetH (residues 2-896) in the absence (blue) and presence of 0.6 mM $CH_3$-$H_4$folate (green). Both the **(A)** resting state and **(B)** homocysteine-methylation state yield reasonable fits to the substrate-free data (blue), while the **(C)** folate-demethylation state does not. Based on the robustness of the full-length $CH_3$-Cob(III) MetH to X-ray exposure, we believe that the resting state (which has a cap-on $B_{12}$ domain) is likely the dominant state in solution in the absence of substrates. By contrast, the **(D)** resting and **(E)** homocysteine states yield poor fits to the $CH_3$-$H_4$folate-incubated data (green), while the **(F)** folate state describes the data well, indicating that the conformational change upon incubation with folate is to favor this conformation. Fits were calculated using CRYSOL[41,42] (Table S11). Model construction is described in the Extended Methods.

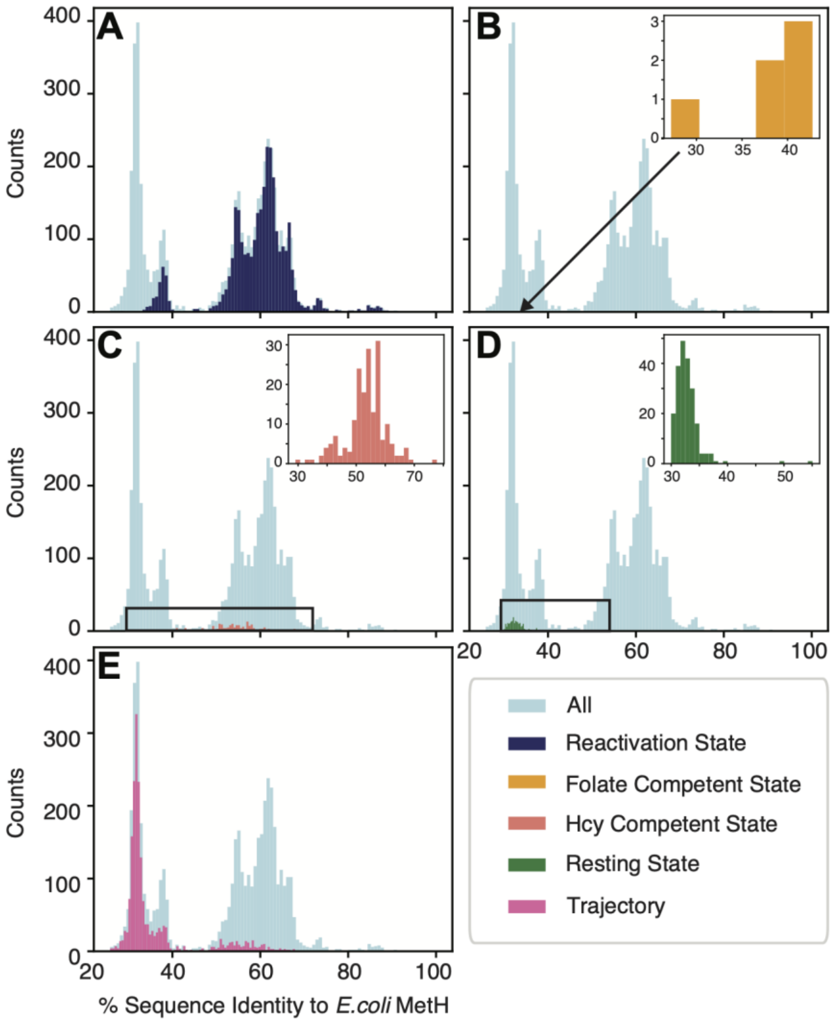


**Figure S20: Analysis of 4-domain MetH sequences in the AlphaFold database by sequence identity to *E.coli* MetH. (A)** MetH sequences predicted to be in the reactivation state (navy blue) have high sequence identity to *E. coli* MetH. **(B)** MetH sequences predicted to be in the folate-demethylation state (orange) have low sequence identity to *E. coli* MetH. Inset shows a zoom in around the 30-40% sequence identity region. **(C)** MetH sequences predicted to be in the Hcy-methylation state (salmon) have moderate sequence identity to *E. coli* MetH. Inset shows a zoom in around the 30-70% sequence identity region. **(D)** MetH sequences predicted to be in the resting state (green) have low sequence identity to *E. coli* MetH. Inset shows a zoom in around the 30-40% sequence identity region. **(E)** MetH sequences predicted to be along the trajectory between resting state and the Hcy-methylation state (pink) as shown in Figure S16 have the lowest sequence identity to *E. coli* MetH. Note that the full histograms have a different binning than the insets.
